# ComFB bound to c-di-GMP reduces swimming speed and tumbling frequency in *B. subtilis*

**DOI:** 10.1101/2025.07.11.664319

**Authors:** Jeanette Hahn, Roberto Noland, Louisa Celma, Abdalla A. Elshereef, Sherihan Samir, Eugenie Dubnau, Khaled A. Selim, Daniel B Kearns, David Dubnau

## Abstract

ComFB is encoded by the *comF* operon in *Bacillus subtilis* together with ComFA and ComFC. Although ComFA and ComFC are essential for natural transformation, ComFB is dispensable, and its role remains unknown. We show here that ComFB binds specifically and with high affinity to the second messenger c-di-GMP, thereby inhibiting swarming over surfaces and swimming through the porous matrix of soft agar by depressing tumbling frequency and reducing swimming speed. While wild-type cells readily navigate the agar matrix, ComFB overexpression causes cells to stall, preventing them from changing direction to escape entrapment in the agar. Suppressors of this deficiency arise from mutations in CheY and in the rotor proteins FliY and FliM, all of which increase tumbling, a phenomenon akin to “pseudotaxis”. These results reveal a c-di-GMP regulatory system that modulates motility independently of MotI, another c-di-GMP receptor that affects motility in *B. subtilis*. ComFB-mediated changes in swimming behavior in the competent state may favor biofilm formation and dispersal to distant localities.

**IMPORTANCE:** Bacterial motility is tightly regulated, and the second messenger c-di-GMP often inhibits flagellar expression and/or function. Revealing the mechanisms of these regulatory pathways is broadly important for understanding bacterial motility and c-di-GMP-related processes, as well as the roles of motility in the transition to biofilm formation. We show that ComFB is a specific, high-affinity receptor for c-di-GMP that modulates flagellar activity in *Bacillus subtilis* by suppressing tumbling and reducing the swimming speed.

## INTRODUCTION

Flagellar motility in liquids and on surfaces is widespread despite the high energetic cost of flagellar synthesis and operation (1). This cost requires that the formation and function of flagella are tightly regulated, often through mechanisms involving the second messenger bis-(3′-5′)-cyclic diguanylic acid (c-di-GMP). c-di-GMP is ubiquitous in bacteria and regulates diverse activities beyond motility, including biofilm formation, exopolysaccharide synthesis, virulence, DNA replication, and cell size (2, 3). To fulfill its regulatory roles, c-di-GMP interacts with receptors, often proteins, which then act on phenotype-specific targets. For example, the *B. subtilis* (*Bsu*) receptor MotI, when bound to c-di-GMP, interacts with the flagellar stator protein MotA to uncouple flagella from the motor force that enables swimming (4).

Recently, a cyanobacterial protein with partial sequence and structural similarity to the *Bacillus subtilis* (*Bsu*) protein ComFB was shown to bind c-di-GMP (5). In cyanobacteria, ComFB regulates type IV pilus-dependent functions, specifically phototactic motility and natural competence, by modulating minor pilin gene expression through interactions with the transcription factors SyCRP1 and SyCRP2 (6). In fact, ComFB*^Bsu^*also binds c-di-GMP (7) with high specificity and affinity, as shown here. While MotI is a PilZ-type c-di-GMP binding receptor with canonical RR and (N/D)hSXXG motifs (8), ComFB represents a new class of cyclic-dimeric nucleoside monophosphate-binding proteins, widespread in bacterial phyla (5, 7, 9). ComFB is embedded in the *comF* operon of *Bsu*, which encodes two additional proteins, ComFA and ComFC, needed for the transport of transforming DNA to the cytoplasm (10–13). In *Bsu*, the *comF* operon is expressed in a minority of cells under control of the transcriptional activator ComK, during the transition to stationary phase (14–18). In the eponymous “K-state”, about 20 genes required for transformation are transcribed, along with about 80 additional genes that are dispensable for transformation and have largely uncharacterized roles. One of these is *comFB* (19–21), which is co-transcribed with *comFA* and *comFC* but is not required for transformation (11), suggesting it plays a K-state role distinct from transformation. Consistent with its dispensability for transformation, Firmicute *comF* operons consistently encode ComFA and ComFC, but only some encode ComFB (8). A search using eggNOG v6 (22) revealed that although 447 species encode orthologs of ComFB*^Bsu^*, these are sporadically distributed, including among more phylogenetically distant bacteria, e.g., cyanobacteria, *Vibrio cholerae*, and *Treponema denticola* (5, 7). Although the atomic structure of *ComFB^Bsu^* has been solved in the apo form (11), its binding to c-di-GMP suggests a regulatory role.

Here, we report that ComFB binds c-di-GMP with high affinity and specificity and that its overexpression inhibits swarming across an agar surface and swimming through soft agar. These inhibitory effects are c-di-GMP-dependent. ComFB overexpression does not prevent swimming in agar-free medium; instead, it reduces swimming velocity and tumbling frequency, causing cells to exhibit persistent forward swimming. Suppressors that arise in the response regulatory protein CheY and in the switch complex proteins FliY and FliM increase the tumbling frequency and restore migration through soft agar, a phenomenon reminiscent of pseudotaxis (23, 24).

## RESULTS

### ComFB binds c-di-GMP specifically and with high affinity

As noted above, orthologs of ComFB have been reported to bind cyclic dinucleotides, including c-di-GMP and c-di-AMP (5, 7, 9). To gain further insight into the specificity and affinity of *ComFB^Bsu^*binding to c-di-GMP using isothermal titration calorimetry (ITC), we first purified it in its native dimeric state (Fig. S1). The raw titration data of c-di-GMP with ComFB were fitted using a single-site binding model to determine the average dissociation constant (KD) for all available binding sites (Table 1, Fig. 1A). ITC experiments revealed that *ComFB* has a high affinity for c-di-GMP (KD ∼140 nM; Table 1), comparable to that of other high-affinity c-di-GMP binders, e.g., PilZ and FimX (25). In contrast, the ComFB KD value for binding to c-di-AMP (Fig. 1B) (9), which *Bsu* also produces (26), is nearly 500-fold higher (Table 1), above the likely physiological concentration.

**Fig. 1.**
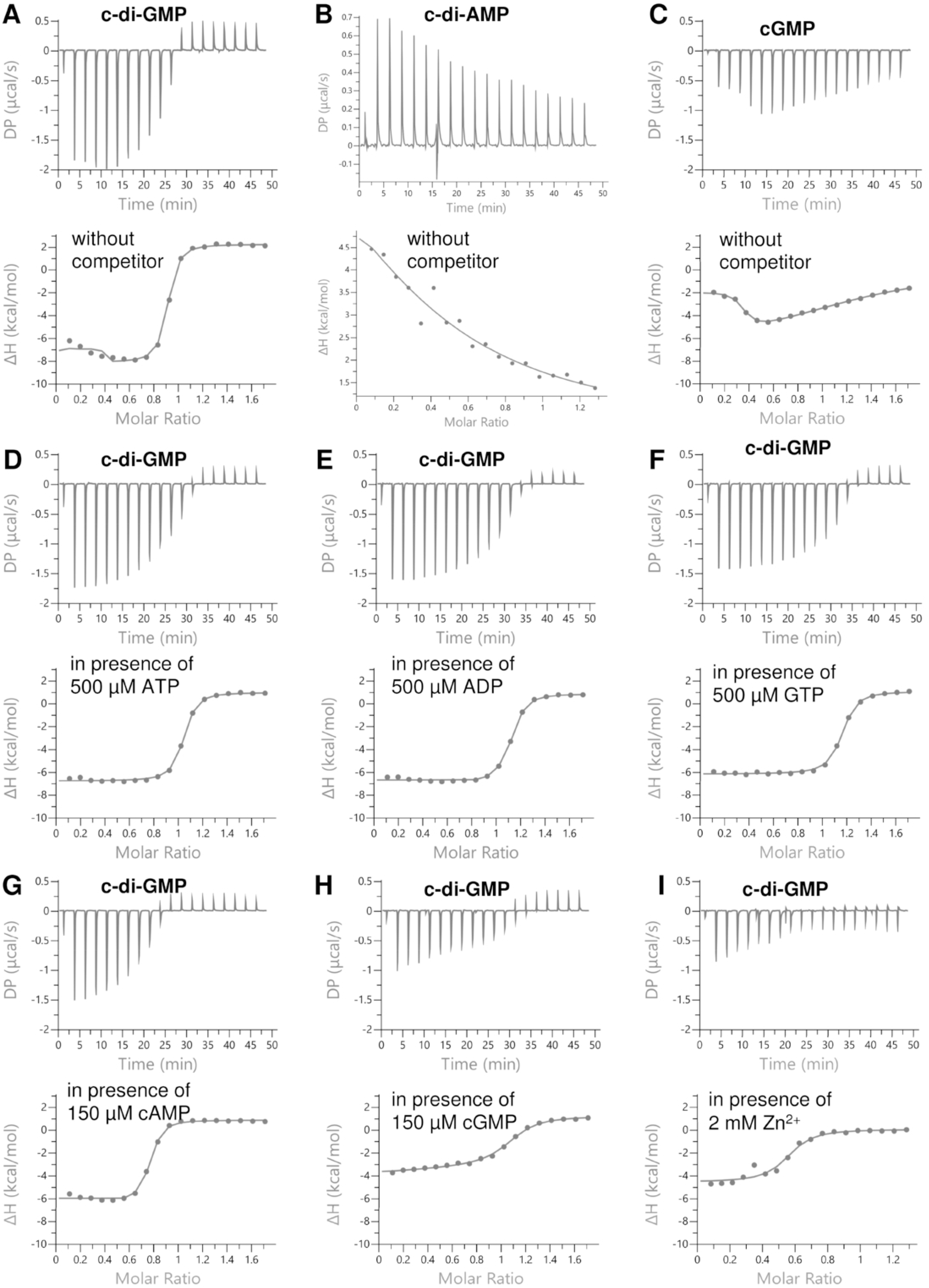
Isothermal titration calorimetry (ITC) analysis of c-di-GMP, cGMP, and c-di-AMP binding to ComFB, as indicated. Upper panels show the raw ITC data in the form of heat produced during the titration of the respective nucleotide on the ComFB protein; lower panels show the binding isotherms and the best-fit curves according to the one-binding-site model. (A-C) ITC analysis of c-di-GMP (A), c-di-AMP (B), and cGMP (C) binding to ComFB in the absence of competitors. (D-I) ITC analysis of c-di-GMP binding to ComFB in the presence of ATP (D), ADP (E), GTP (F), cAMP (G), cGMP (H), or Zn^2+^ (I).

**Table 1.**
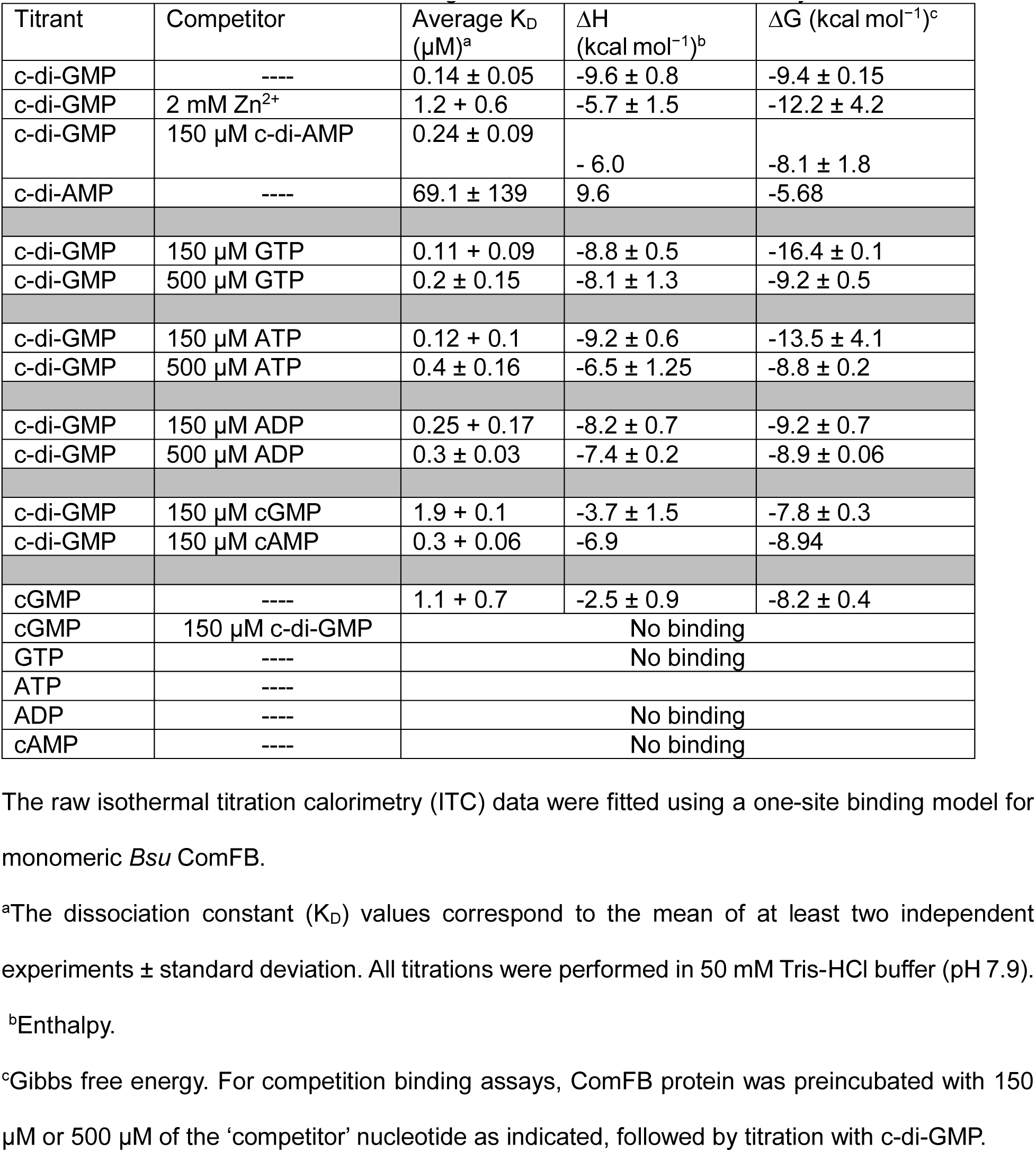
c-di-GMP binding to BusComFB as determined by ITC.

Accordingly, c-di-GMP competes efficiently with c-di-AMP at a saturating concentration of 150 µM (Table 1).

To further explore the specificity of c-di-GMP binding, we tested other, more abundant nucleotides (ATP, ADP, and GTP) and the nucleotides cGMP and cAMP, the latter two of which have not been reported to be present in *Bsu*. GTP, ATP, and ADP could not bind to ComFB (Fig. S2), and their presence at 150 or 500 µM did not significantly affect the c-di-GMP affinity to ComFB (Table 1, Figs. 1D-E and S2). However, at the higher competitor concentration of 500 µM, the binding enthalpy of c-di-GMP was slightly reduced (Table 1; compare isotherm of Fig. 1A with 1D-F), causing a slight increase in the KD values (Table 1). cAMP could not bind ComFB (Fig. S2G), but its presence at 150 µM triggered a trivial decrease in the c-di-GMP binding enthalpy to ComFB (Table 1), yielding weaker isotherms (compare the isotherm of Fig. 1A with 1G). We conclude that among the nucleotides tested so far, only c-di-GMP interacts significantly with ComFB.

In contrast, cGMP bound ComFB with a KD of ∼1.1 µM. The cGMP titration curve showed a complex biphasic shape; in the first injections, the enthalpy increased until a maximum was reached at a cGMP concentration of ∼40 µM (Fig. 1C). Subsequently, the heat isotherm gradually decreased, indicating gradual saturation of ComFB by cGMP (Fig. 1C). The binding of cGMP was completely abolished in the presence of a saturating concentration of c-di-GMP (150 µM) (Fig. S2H). In the presence of 150 µM cGMP, c-di-GMP was able to compete efficiently, replacing cGMP for ComFB binding, although the apparent KD for c-di-GMP binding increased by more than 10-fold in the presence of cGMP (Table 1), as indicated by a decrease in the binding enthalpy (Table 1; compare isotherm of Fig. 1A with 1H). Thus, cGMP is unable to efficiently occupy the binding sites of ComFB. cGMP is produced by some bacteria (9, 27), but to our knowledge, it has never been reported to exist in the Firmicutes, and its weak interaction with ComFB is likely not biologically significant. We conclude that ComFB is a c-di-GMP receptor in *Bsu* that binds c-di-GMP with high affinity and specificity; it does not interact with other purine nucleotides present in *Bsu*.

### ComFB inhibits swarming and swimming through agar when bound to c-di-GMP

c-di-GMP binding receptors have been implicated in inhibiting swimming and swarming motility in several bacterial species (4, 28, 29). Swarming, bacterial expansion across a solid surface, depends in *Bsu* on the association of individual cells into “rafts”, on the secretion of surfactin, and on the SwrA protein, which induces hyperflagellation. To study swarming, the undomesticated NCIB 3610 background is used, because domesticated strains do not produce SwrA or surfactin (30). Swimming, measured as cell movement through soft agar, is independent of these factors and so can be studied in domesticated *Bsu* strains.

To determine whether the overexpression of ComFB affected swarming and swimming, we expressed ComFB from the *amyE* locus under the control of a constitutive promoter (Pc) (31), using either a weak or a strong ribosomal binding site (RBS). The “weak” construct (*amyE::*Pc(w)*comFB*) employed the native ComFB RBS, while the “strong” construct (*amyE::*Pc(s)*comFB*) utilized the RBS of the highly expressed *gapA* gene. In all strains, the native *comFB* remained intact but was expressed at a low level, as the experiments were conducted in LB agar, in which the K-state is not activated (see Fig. 2A). To test the dependency of ComFB on c-di-GMP signaling, each construct was tested in the wild-type background and in the *pdeH* deletion background. PdeH is the only known c-di-GMP phosphodiesterase in *Bsu,* and in its absence, the cellular concentration of c-di-GMP increases (31, 32). Western blotting showed that *comFB* expression from the weak construct slightly augmented the total amount of ComFB, while the strong construct increased ComFB about 100-fold (Fig. 2A). Importantly, the expression of ComFB from the strong overproduction construct was not affected when *pdeH* was deleted. Although the *amyE*::Pc(w)*comFB* construct did not affect swarming on its own, it exhibited an inhibitory effect when combined with *ΔpdeH* (Fig. 2B). As reported previously (32), *ΔpdeH* itself depressed swarming, but the inhibition was enhanced when combined with *amyE*::Pc(w)*comFB*. In contrast, the *amyE::*PC(s)*comFB* construct partially inhibited swarming on its own and eliminated it when *pdeH* was deleted (Fig. 2C). These results are consistent with the conclusion that ComFB inhibits swarming when bound to c-di-GMP.

**Figure 2.**
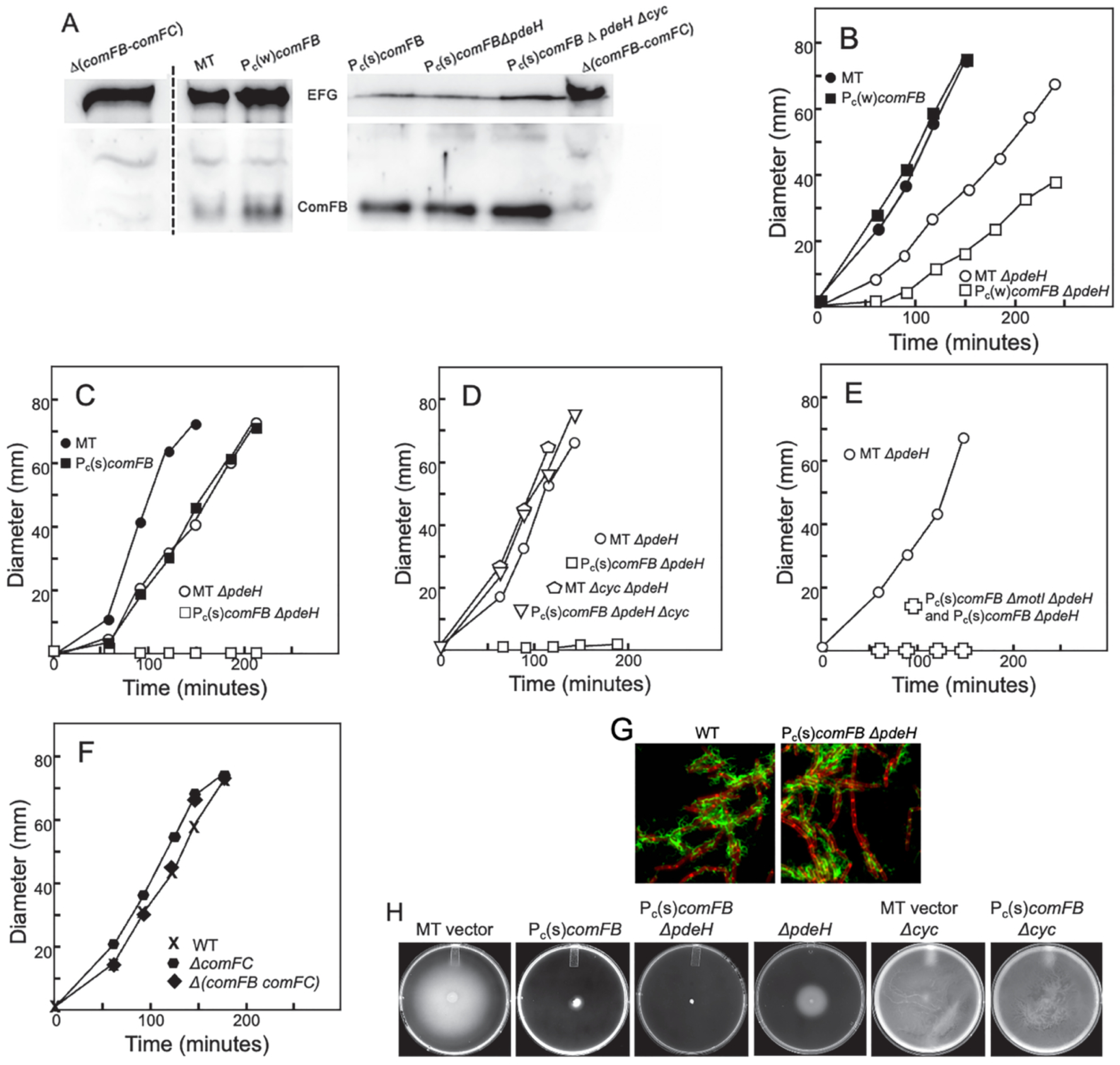
ComFB overexpression, together with c-di-GMP, inhibits swarming and swimming in agar. To avoid crowding, FB and FC were used as abbreviations for ComFB and ComFC. (A) Overexpression of the weak and strong versions of *comFB*. Western blots were performed with anti-ComFB antibodies. As a loading control, the tops of the membranes were separated and then developed with anti-EFG (elongation factor G) antibodies. The Δ(*FB-FC*) strain confirms the identity of the ComFB signal. The vertical dashed line indicates that intervening lanes were omitted from a gel. For the gel on the right, the first three lanes were loaded with a 20-fold-diluted lysate (2 μl), whereas the other lanes contained 15 μl of undiluted lysate. *Δcyc* denotes strains deleted for the three diguanylate cyclases of *Bsu* (*dgcP*, *dgcK*, *dgcW*). The constructs *amyE::*Pc(w)*comFB* and *amyE*::Pc(s)*comFB* are described in the text. MT is the empty vector strain with pKB149 (31) in *amyE*. The MT vector strain produces a low level of ComFB, which is increased moderately in the *amyE*::Pc(w)*FB* strain and by about 100-fold in the *amyE*::Pc(s)*comFB* strain. (B-F) Each panel exemplifies at least three repeated swarming experiments on 0.7% LB agar plates. The cell sheet expansion diameters were measured at intervals and plotted against time. The symbols are consistent across all five panels, with empty symbols indicating *ΔpdeH* strains. (G) Swimming in soft (0.3%) agar of the indicated strains. (H) Flagellar staining, in which the indicated strains, both carrying the *hag^T209C^* allele, were reacted with Alexa Fluor 488 C5 maleimide (green), stained with the membrane dye FM4-64 (red), and imaged as described in the Materials and Methods section.

To further investigate the role of c-di-GMP, we used the *amyE::*Pc(s)*FB* construct in a c-di-GMP null strain (*Δcyc*), lacking the three diguanylate cyclases of *Bsu* (*dgcW*, *dgcP,* and *dgcK*) (31). In this background, the strong overproduction of ComFB did not affect swarming (Fig. 2D), and Western blotting showed that the absence of the cyclases did not reduce the amount of ComFB (Fig. 2A). These results strongly support a dose-dependent inhibition of swarming by ComFB when bound to c-di-GMP.

Next, we sought to determine whether the swarming inhibition by ComFB requires the candidate mediators, MotI and ComFC. MotI is a c-di-GMP receptor that prevents motility by disengaging the flagellar stator protein MotA (4). To determine whether MotI mediates the inhibitory effect of ComFB, we placed the *amyE::*Pc(s)*comFB* construct in a *ΔmotI ΔpdeH* background. As shown in Fig. 2E, the inhibitory effect of ComFB on swarming is not reversed when *motI* is deleted. Thus, *Bsu* encodes two c-di-GMP receptors that independently inhibit swarming. ComFC is a protein co-expressed with the gene encoding ComFB. To explore whether ComFB and ComFC might be functionally related, we used a markerless construct in which nearly the entire *comFC* coding sequence was removed. This construct was derived from a *ΔcomFC::erm* deletion by removing the Erm-cassette using plasmid pDR244 (33). This was necessary because the cassette carries a downward-facing promoter, and readthrough from the ComK-dependent promoter has been shown to enhance FlgM production, leading to the loss of flagella (34). The deletion of *comFC* had no detectable effect on swarming (Fig. 2F), nor did the deletion of both *comFB* and *comFC* genes, consistent with weak *comFB* expression in LB media, where the K-state is not expressed.

The *amyE*::Pc(s)*comFB* construct also strongly inhibited the expansion of domesticated *Bsu* through 0.3% soft agar, even without the inactivation of *pdeH*, although more complete inhibition was observed when *pdeH* was deleted (Fig. 2G). When the three diguanylate cyclases were deleted in the *amyE*::Pc(s)*comFB* strain, movement through agar was restored (2H).

One way the ComFB:c-di-GMP complex might inhibit swarming and swimming is by interfering with flagella formation. To monitor flagellar abundance, cells carrying a *hagT209C* mutation were stained with maleimide-Alexa Fluor 488 (35). Straining revealed that the strong overproduction of ComFB does not obviously interfere with flagellar production even when *pdeH* was deleted (Fig. 2H), although this result does not exclude subtle effects on the number of flagella. We conclude that ComFB likely interferes with both swarming and swimming through agar by impacting flagellar activity, rather than inhibiting flagellar biosynthesis, surfactin, or SwrA.

### Suppressors of ComFB overexpression

To gain insight into the ability of ComFB to inhibit swimming through agar, and with the hope that we might identify a ComFB target, we selected suppressors that restored swimming ability, using a domesticated parent strain with the genotype *ΔpdeH amyE::*Pc(s)*comFB thrC::*Pc(s)*FB*. Two copies of Pc(s)*comFB* were included to minimize the chance of isolating ComFB loss-of-function mutations, and the *ΔpdeH* allele was included to enhance the inhibitory effect. Individual colonies were stabbed into LB plates containing 0.3% agar. After 72 hours of incubation at 37 °C, nearly every plate contained a flare emerging from the centrally located colony, suggesting that suppressors arose in one or more genes. Cells were isolated from 15 of the flares and were submitted for whole-genome sequencing. Mutations were identified in the histidine kinase *cheA,*the response regulator *cheY*, and in the rotor genes *fliM, and fliY*. Re-testing showed that the *cheA* mutations, all of which were deletions, frameshifts, or stop codons, exhibited only minimal, inconsistent suppression and were not studied further. The identities of the *cheY*, *fliY,* and *fliM* suppressor strains are shown in Table 2.

**Table 2.** Suppressor Mutations.

| Gene | Mutation | Position<sup>1</sup> | Effect |
| --- | --- | --- | --- |
| *cheY* | CCA->CTA | 173 | P58L |
| *cheY* | ATG->ATA | 267 | M89I |
| *cheY* | GAT->TAT | 274 | D92Y |
| *cheY* | GCA->GTA | 341 | A114Y |
| *fliM* | ACT->CCT | 205 | T69P |
| *fliY* | ACT->CCT | 202 | T68P |
| *fliY* | GCG->GTG | 335 | A111V |
| *fliY* | TCT->TTT | 575 | S192F |
| *fliY* | ATA->ACA | 602 | I201T |

<sup>1</sup>Position in the coding sequence

To confirm the suppressors, several of those in *cheY* and *fliY* were swapped into wild-type and *amyE::*Pc(s)*comFB* backgrounds, without the deletion of *pdeH*, to restrict effects to ComFB overexpression. In all cases, the mutations restored mobility through soft agar to the *amyE::*Pc(s)*comFB* strain, even causing more rapid expansion than with the wild-type parent (Fig. 3). Without ComFB overexpression, the mutations also caused more rapid expansion than that of the wild-type strain. Viewed in wet mounts, the *cheY, fliY, and fliM* suppressor mutations in the absence of overexpressed ComFB obviously increased the frequency of tumbling (see Movies S1 and S2 as examples), compared to the wild-type (Movie S3). Cells of all the suppressor mutant strains exhibited frequent tumbles, interspersed by relatively short runs. It has been reported that the failure of *E. coli* and *A. tumefaciens* mutants with run biases to navigate through soft agar was similarly suppressed by mutations that increased tumbling (23, 24), a phenomenon known as pseudotaxis. This suggested that ComFB overexpression may cause a run bias. Although FliY and FliM are not excluded as potential ComFB targets, our suppressor mutations did not appear to act as target mutations but rather bypassed the effect of ComFB by causing an increase in tumbling.

**Fig. 3.**
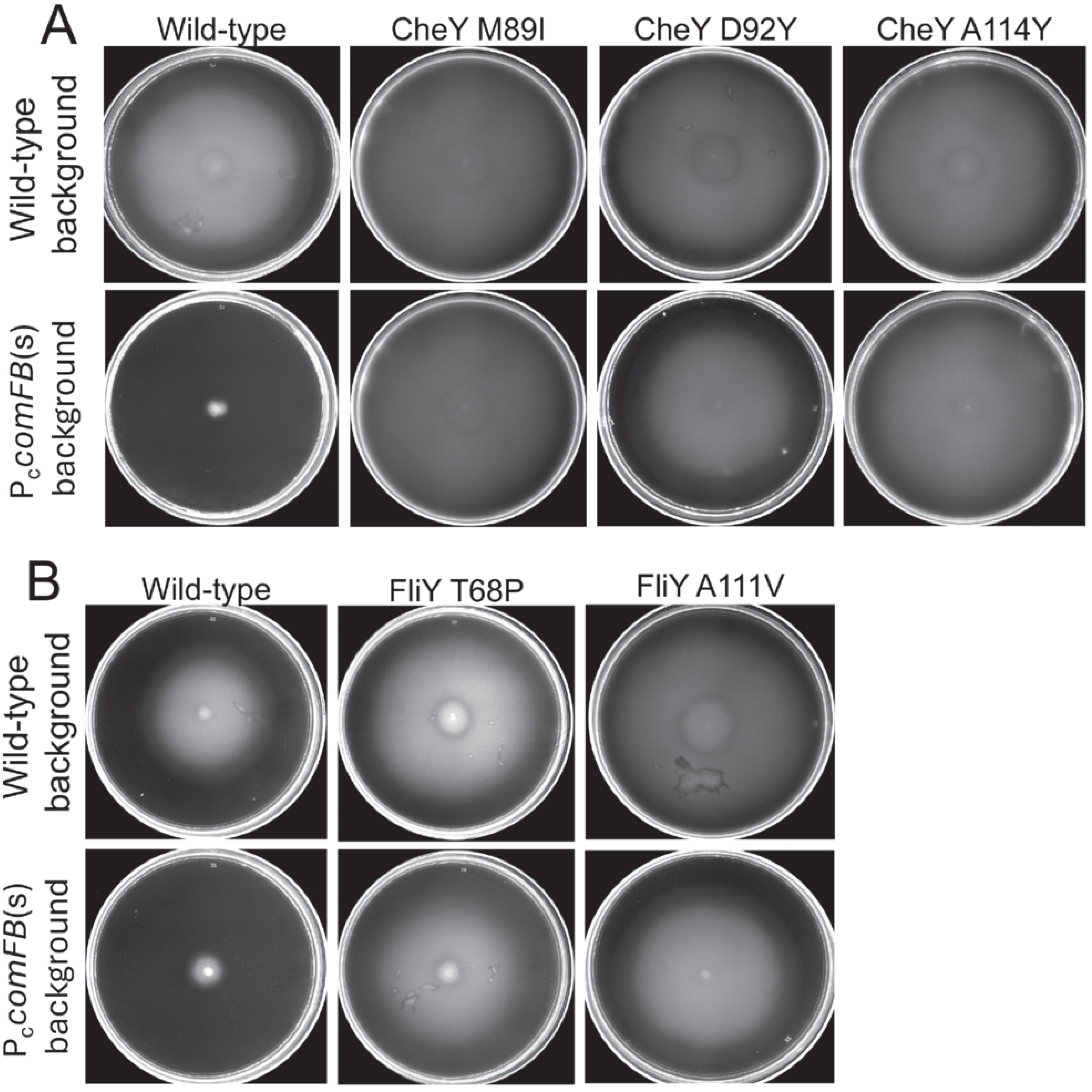
Movement through soft agar of strains bearing the indicated suppressor mutations in the wild-type and Pc(s)*comFB* backgrounds. (A) Suppressor mutations in *cheY.* (B) Suppressor mutations in *fliY*

### The effect of ComFB overexpression on swimming behavior

To further characterize the ComFB-associated motility defect, we microscopically examined the swimming behavior of the wild-type (Movie S3) and the *amyE*::Pc(s)*comFB* strain (Movie S4). Despite its near-complete inability to navigate through LB agar (Fig. 2), the *amyE*::Pc(s)*comFB* strain swam actively in agar-free LB, exhibiting long runs. One subtle difference in the wild-type and ComFB overexpressing strains is visible in these videos. In the wild-type (Movie S3), an occasional cell abruptly dives out of the field of view, indicating a tumbling event. In contrast, few if any such events were recorded for the ComFB overexpression strain (Movie S4). When movment though 0.3% agar was viewed under the microscope, wild-type cells exhibited runs punctuated by brief pauses, during which they idled in place before resuming another short run in a different direction (Movie S5). This behavior qualitatively resembles the swimming of *E. coli* in soft agar, as described by Wolfe and Berg (24), and the “hopping and trapping” motility observed during swimming in synthetic porous media (36). In contrast, ComFB overexpression had a dramatic effect on movement in agar. Many cells remained idle, sometimes quivering in place and unable to move through the agar matrix, while a few cells were able to navigate what are likely open channels in the agar (Movie S6). The latter behavior resembles that of *E. coli* and *Agrobacterium tumefaciens* mutants with reduced tumble frequencies swimming in soft agar (23, 24).

For a quantitative description of how ComFB affects swimming dynamics, we used MiMoSA (37), a modified version of RABiTPy (38), following the analytic framework described by Najafi et al (39). Table 3 and Fig. 4 show that the run (VR) speed is reduced in the ComFB overexpression strain, whereas the translational tumble speed (VT) is unaffected. Additionally, the overexpression strain has a reduced tumble bias, computed in two slightly different ways (Table 3). When it tumbles, it exhibits a reduced average turning angle, reflected in a larger value of R, defined as cosΦ, where Φ is the angular change in direction between successive runs (39). The increased value of R (Table 3) and the reduced frequency of detected tumbles indicate that ComFB overexpression increases the likelihood that a cell will continue moving in the same direction. The relatively small standard deviation for R in the overexpression strain, despite its smaller sample size, is consistent with the existence of a restraint in its directional change between runs, compared with the wild-type strain, which tumbles more freely. The nonzero value of VT confirms that cell translation continues during a tumble, albeit more slowly than during a run (39), suggesting that incomplete disassembly of flagellar bundles may sometimes occur. Together, our findings show that ComFB overexpression causes cells to swim with a reduced probability of changing direction and at a slower speed than wild-type cells. This explains why ComFB interferes with movement through a porous medium, where cells often become trapped in narrow spaces. Escape from such cul-de-sacs requires reorientation (tumbling), which is inhibited by ComFB, consistent with genetic suppression by increased tumbling.

**Fig. 4.**
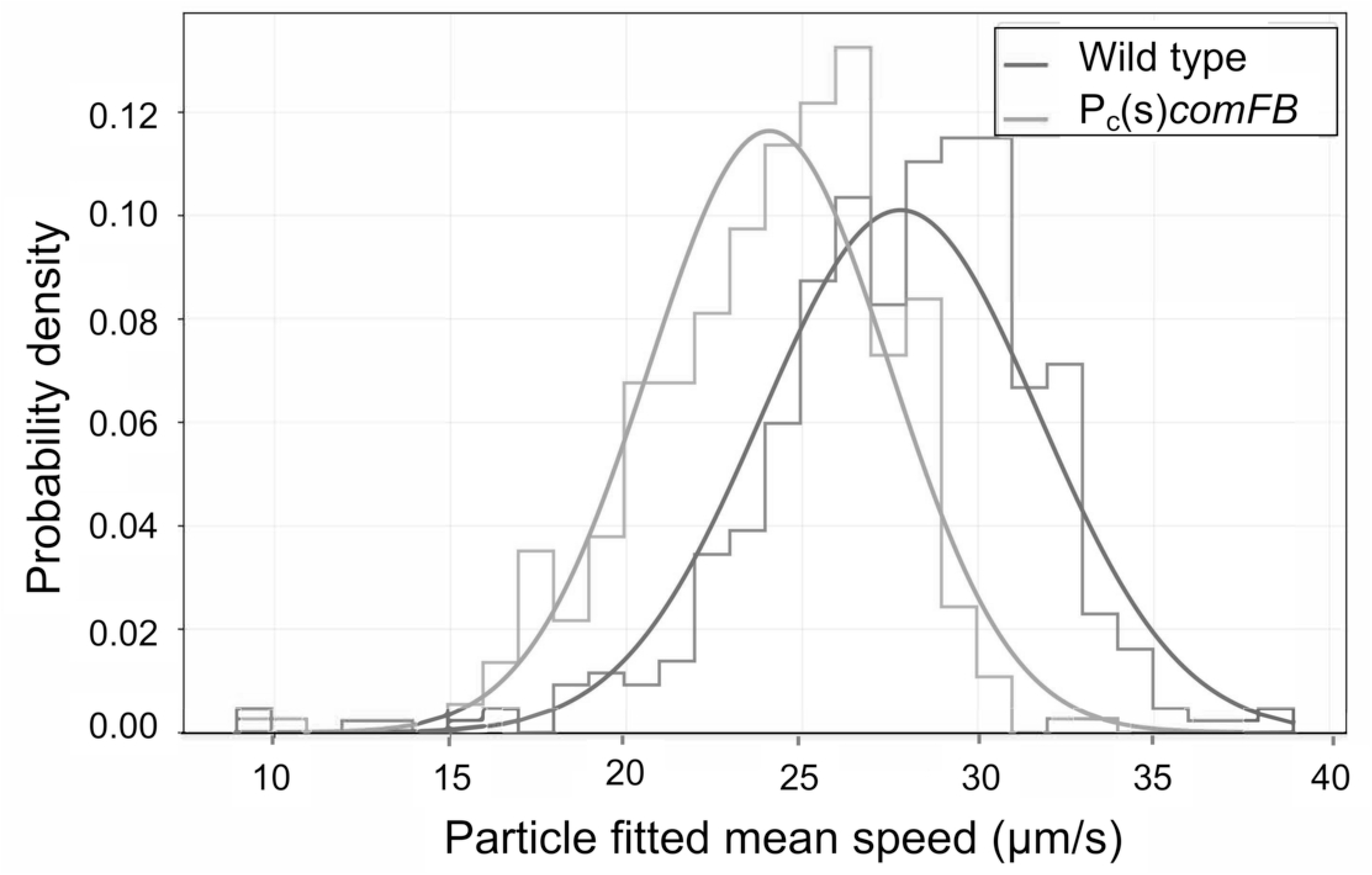
Fitted speed distributions of Wild-type (blue) and *amyE*::Pc(s)*comFB*) (orange) strains. The curves were fitted to the binned observational data at a 95% confidence interval. The mean values for the wild type and ComFB overexpression strains were 27.8 μm/s and 24.1 μm/s, with n=434 and 369, respectively.

**Table 3.**
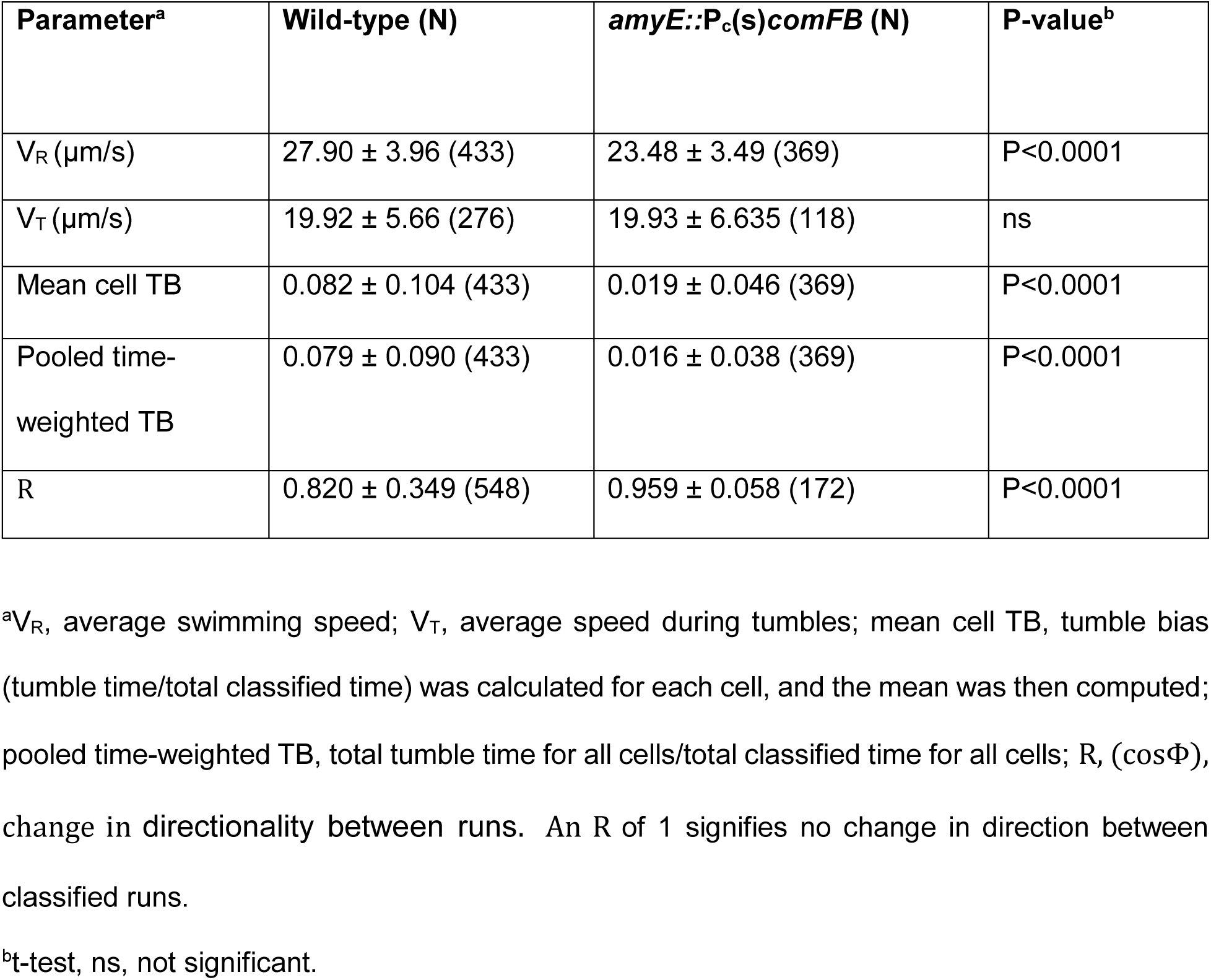
Swimming statistics.

| Parameter<sup>a</sup> | Wild-type (N) | *amyE::P<sub>c</sub>(s)comFB* (N) | P-value<sup>b</sup> |
| --- | --- | --- | --- |
| V<sub>R</sub> (μm/s) | 27.90 ± 3.96 (433) | 23.48 ± 3.49 (369) | P<0.0001 |
| V<sub>T</sub> (μm/s) | 19.92 ± 5.66 (276) | 19.93 ± 6.635 (118) | ns |
| Mean cell TB | 0.082 ± 0.104 (433) | 0.019 ± 0.046 (369) | P<0.0001 |
| Pooled time-weighted TB | 0.079 ± 0.090 (433) | 0.016 ± 0.038 (369) | P<0.0001 |
| R | 0.820 ± 0.349 (548) | 0.959 ± 0.058 (172) | P<0.0001 |

<sup>a</sup>V<sub>R</sub>, average swimming speed; V<sub>T</sub>, average speed during tumbles; mean cell TB, tumble bias (tumble time/total classified time) was calculated for each cell, and the mean was then computed; pooled time-weighted TB, total tumble time for all cells/total classified time for all cells; R, (cosΦ), change in directionality between runs. An R of 1 signifies no change in direction between classified runs.
<sup>b</sup>t-test, ns, not significant.

### FliY T68P and A111V neither prevent swarming nor suppress inhibition by **amyE::Pc(s)comFB**

The suppression of tumbling and the increased persistence imposed by ComFB do not readily explain how ComFB overexpression inhibits swarming, because swarming of *B. subtilis* and other bacteria is typically biased toward runs (40, 41). To shed light on this, we examined the effects of FliY T68P and A111V on swarming in the wild-type 3610 background and tested if these mutations suppressed the absence of swarming of the *amyE*:: *Pc(s)*comFB *ΔpdeH* strain. Fig. 5 shows that whereas FliY A111V had no discernible effect on swarming, the FliY T68P strain began to swarm, but its rate of expansion gradually decreased. Cells taken from the advancing edge of these swarms confirmed that the tumble bias of the mutants was retained in swarming cells. The FliYT68P strain is shown as an example (Movie S7), compared with cells from the wild-type swarm cells (Movie S8). showing that the mechanism that suppresses tumbling during swarming (40, 41) is not epistatic on the *fliY* mutations. Even with their tumble bias, these mutations evidently allow sufficient cell translation for swarming to occur, consistent with the nonzero VT value noted above and in Table 3. Significantly, neither *fliY* mutation bypassed the inhibition of swarming by ComFB overexpression, implying that the inhibition was not due to decreased tumbling, suggesting that ComFB affects swarming by reducing flagellar motor output (Fig. 4, Table 3).

**Fig. 5.**
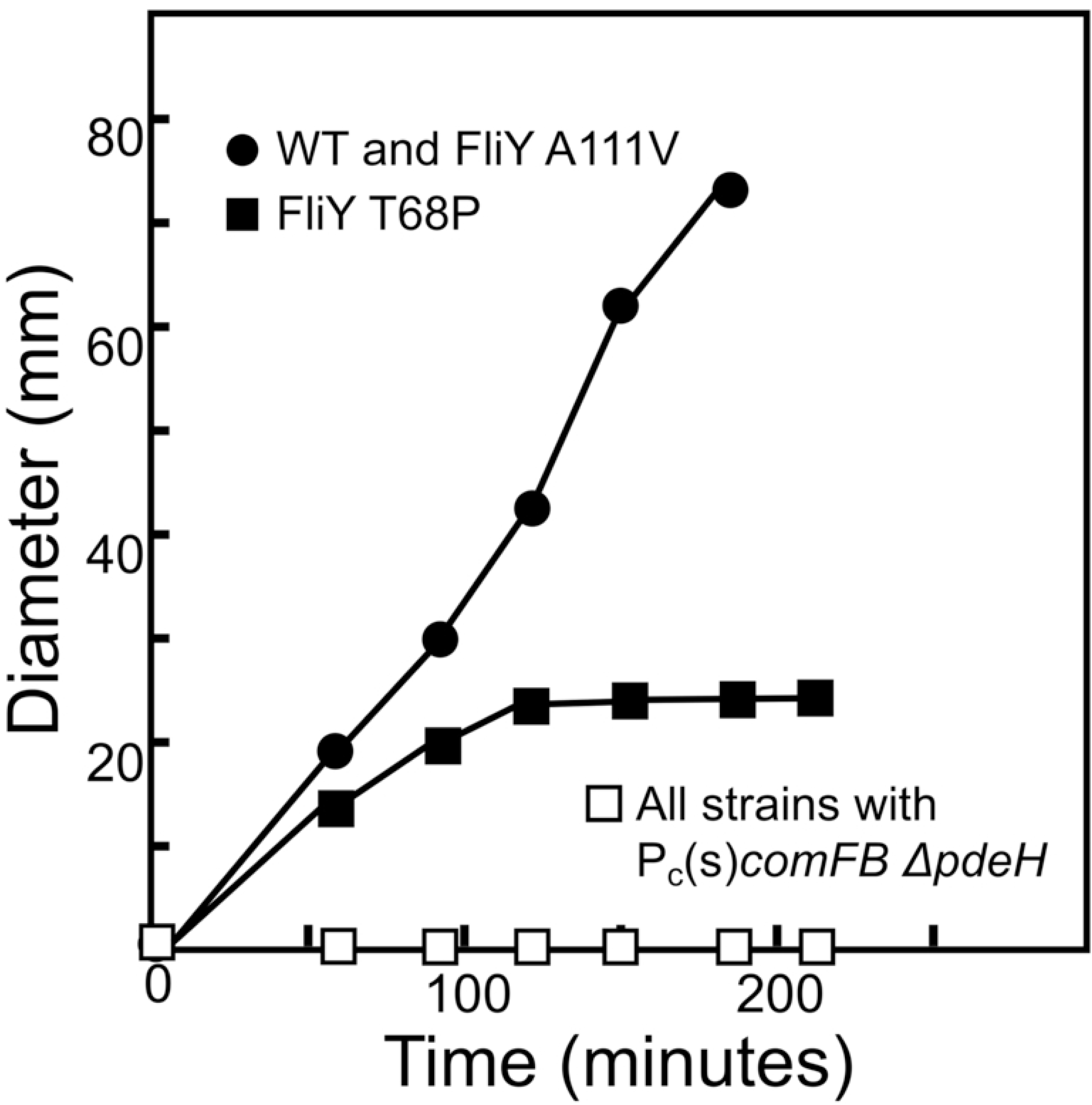
Effects of FliY mutations on swarming in the wild-type and *amyE*::Pc(s)*comFB* Δ*pdeH* backgrounds. The *amyE*::Pc(s)*comFB* Δ*pdeH* strains did not swarm, even with the *fliY* mutations.

## DISCUSSION

ComFB represents a new class of c-di-GMP receptors (7, 9), joining three others in *Bsu* that regulate aspects of cell physiology; MotI inhibits motility (4), while YdaK and YkuI have been proposed to regulate the production of uncharacterized extracellular polysaccharides (42, 43). These four receptors belong to distinct protein families; YdaK is a degenerate GGDEF protein lacking cyclase activity, YkuI is an EAL domain protein lacking phosphodiesterase activity, MotI is a PilZ domain protein (31, 32), and ComFB is the founding member of a novel class of receptors (7).

ComFB reduces swimming speed and suppresses tumbling without causing a noticeable decrease in flagellar abundance (Figs. 2H, 4, and Table 3). For these effects, ComFB must be bound to c-di-GMP: the inactivation of *pdeH* augments ComFB’s activity (Fig. 2B, C, and G); ComFB does not inhibit swarming or swimming through agar in a diguanylate cyclase null background (Fig. 2D and G); and it is a specific and high-affinity c-di-GMP binding protein ((7) and Table 1). Swimming in agar requires bacteria to navigate through pores in a maze of agar polymers, a process that is clearly impaired when ComFB is overexpressed. It has been proposed that tumbling is required for bacteria to escape trapping in agar matrix cul-de-sacs (23, 24), and our data show that mutations in CheY, FliM, and FliY that increase tumbling also suppress the soft agar swimming deficiency imposed by ComFB. FliY mutations do not suppress the inhibitory effect of ComFB on swarming, nor do they completely abrogate swarming in the wild-type background (Fig. 5). ComFB may inhibit swarming not by suppressing tumbling, but rather by reducing swimming speed, perhaps similar to that observed when swarming was abolished by disruption of the torque-enhancing protein, SwrD (60). Similarly, ComFB was recently shown to inhibit phototactic motility in cyanobacteria in a c-di-GMP-dependent manner by regulating type IV pili, as the Δ*comFB* mutant exhibited enhanced phototactic motility compared to wild-type cells (6). This suggests that ComFB possesses a conserved function, regulating diverse bacterial motility machines in phylogenetically distant bacteria (7), likely through distinct mechanisms.

In addition to ComFB, *Bsu* encodes EpsE and MotI, additional regulators of flagellar function (4, 44). EpsE contributes to the synthesis of a matrix polysaccharide and is transcriptionally activated under biofilm control. In addition to its biosynthetic activity, EpsE, which does not seem to bind c-di-GMP, acts as a molecular clutch, most likely by binding to the rotor protein FliG and disrupting its interaction with MotA (35, 44). The two-domain protein MotI also acts as a clutch by decoupling the stator protein MotA from the rotor (4). The N-terminal domain of MotI resembles the PilZ domain of YcgR, a well-studied, multi-domain c-di-GMP-binding receptor. Like ComFB, YcgR reduces swimming speed and biases motility toward runs. It does so in enteric bacteria by associating with FliG, FliM, and MotA (45–48). Similar to YcgR, MotI, and EpsE, ComFB is likely to affect rotor mechanics because suppressors that bypass its inhibitory effect were mapped to chemotaxis proteins and to components of the C-ring that control directional changes. However, this hypothesis requires further investigation and should be regarded with caution. Najafi et al. (39) have shown that reducing the number of *B. subtilis* flagella over a 4-fold range lowers the tumbling frequency, albeit without affecting swimming velocity during runs or tumbles, in contrast to our observations with ComFB overexpression (Table 3, Fig. 4).

YcgR and related proteins are considered to favor transitions to a sessile state (45, 46, 49, 50). EpsE, expressed under biofilm control, clearly does the same for *B. subtilis,* and MotI may act similarly. We expect ComFB to be highly expressed in the K-state, and the reduced swimming speed and extended runs imposed by ComFB may favor interactions with surfaces, promoting biofilm formation, although why biofilm formation would be coupled to the K-state is unanswered. EpsE clearly acts when cells are already committed to biofilm formation, and, as a clutch protein, it prevents directed cell movement. Perhaps ComFB acts in K-state cells to promote biofilm initiation, while EpsE subsequently halts motility as cells build a biofilm. Because ComFB alters swimming dynamics to favor continued movement in a given direction, it may promote exploration of distant spaces, increasing the likelihood of encountering transforming DNA or locating nutrients, thereby enhancing the fitness advantage conferred by the K-state.

Two reports document start sites corresponding to a promoter just upstream of *comFB*, both based on data from LB-grown cells. A tiled microarray study of global transcription (50), with a resolution of 22 bases, showed increased transcription 32 bases upstream of the *comFB* open reading frame (position 3,642,139 in the *Bsu* genome). Similarly, Rend-seq (https://rendseq.org/) confirmed the presence of a nearby transcriptional start site at chromosomal position 3,642,135 with single-nucleotide resolution. Fig. 6 shows that the *comFB* and *comFC* coding sequences overlap, suggesting that they are translationally coupled, and that *comFB* is preceded by a plausible SigA promoter sequence located at an appropriate distance upstream from the initiating A residue. Consistent with this, Fig. 2A documents the low-level production of ComFB during growth in LB. These considerations suggest that ComFB, and possibly ComFC, may play a role independent of the K-state.

**Figure 6.**
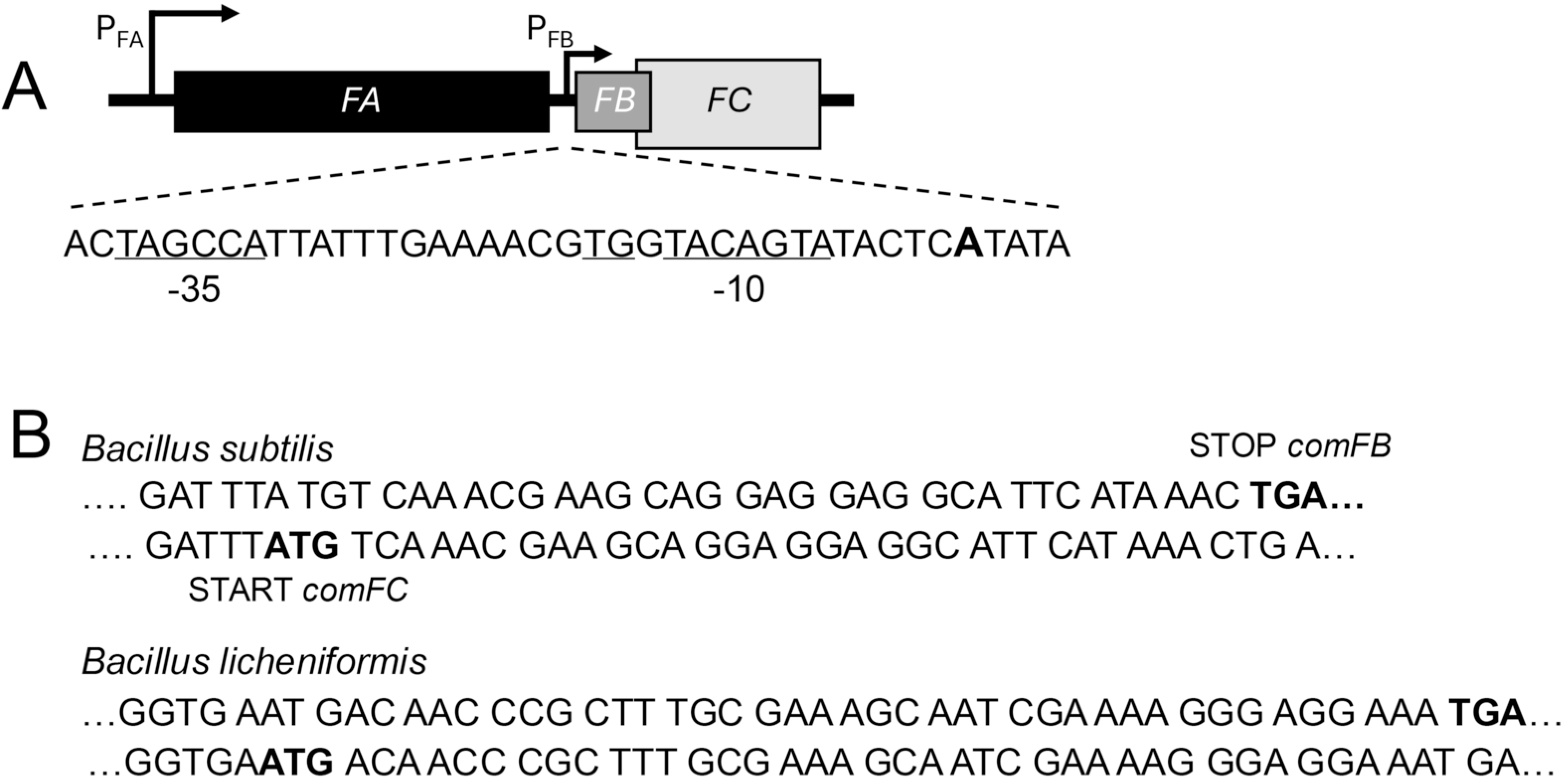
Organization of the comF operon. (A) In Bsu, the comFA, comFB, and comFC open reading frames are transcribed from PFA, a ComK-dependent promoter. PFB, a ComK-independent promoter, drives comFB and comFC, which overlap. The existence of PFB is documented as discussed in the Discussion. The +1 A-residue for PFB is shown in boldface, and the proposed PFB promoter sequence features are underlined. (B) Two examples of bacteria that encode ComFB in their comF operons, demonstrating the overlap of their comFB and comFC coding sequences.

It has been shown that viscous drag on flagella favors transitions to the K-state by downregulating the phosphorylation of the response regulator DegU (51, 52). Interestingly, the *comF* operon is located in the vicinity of the flagella *flgMNKL* operon, placing *comFC* upstream of *flgM*, which encodes an anti-sigma factor for SigD (53), a genomic organization that further supports the involvement of the ComFB protein in regulating flagellar motility. Thus, read-through from the ComK-dependent promoter inhibits flagellar synthesis in K-state cells (34). This suggests an intriguing pattern of regulation: flagellar activity serves as a checkpoint for *entry* into the K-state; further flagellar *synthesis* is restricted in K-state cells; and K-state flagellar *function* may be modulated by ComFB. The connection of ComFB to both motility and competence may extend beyond *Bacillus*: a cyanobacterial ComFB ortholog binds c-di-NMP to regulate both natural competence and pilus biogenesis (9). Because pili are required for twitching motility, these findings underscore the relationships between ComFB, transformation, and motility, albeit with different forms of bacterial locomotion.

## MATERIALS AND METHODS

### Strains, strain construction, and growth conditions

Growth was in LB Medium, except that cells for transformation were grown as described (54). Antibiotics were added to solid media at the following concentrations: chloramphenicol (7 μg/ml), erythromycin and kanamycin (5 μg/ml), spectinomycin (100 μg/ml), and tetracycline (20 μg/ml). Two backgrounds were used as indicated: the domesticated parental strain (IS75) is a derivative of strain 168 and is auxotrophic for histidine, leucine, and methionine. The parental undomesticated strain (BD5776) is a *comI* mutant derivative of NCIB 3610. Constructs were generally introduced by transformation into both backgrounds, but in some cases by transduction with bacteriophage SPP1. To move suppressor mutations into new backgrounds, ∼2 kbp fragments containing the mutations at their midpoint were generated by PCR and cloned into the pMINIMAD plasmid, which was then used to place each mutation at its native locus, with loss of the plasmid (55). Strains are listed in Table 4 and plasmids in Table 5.

**Table 4.**
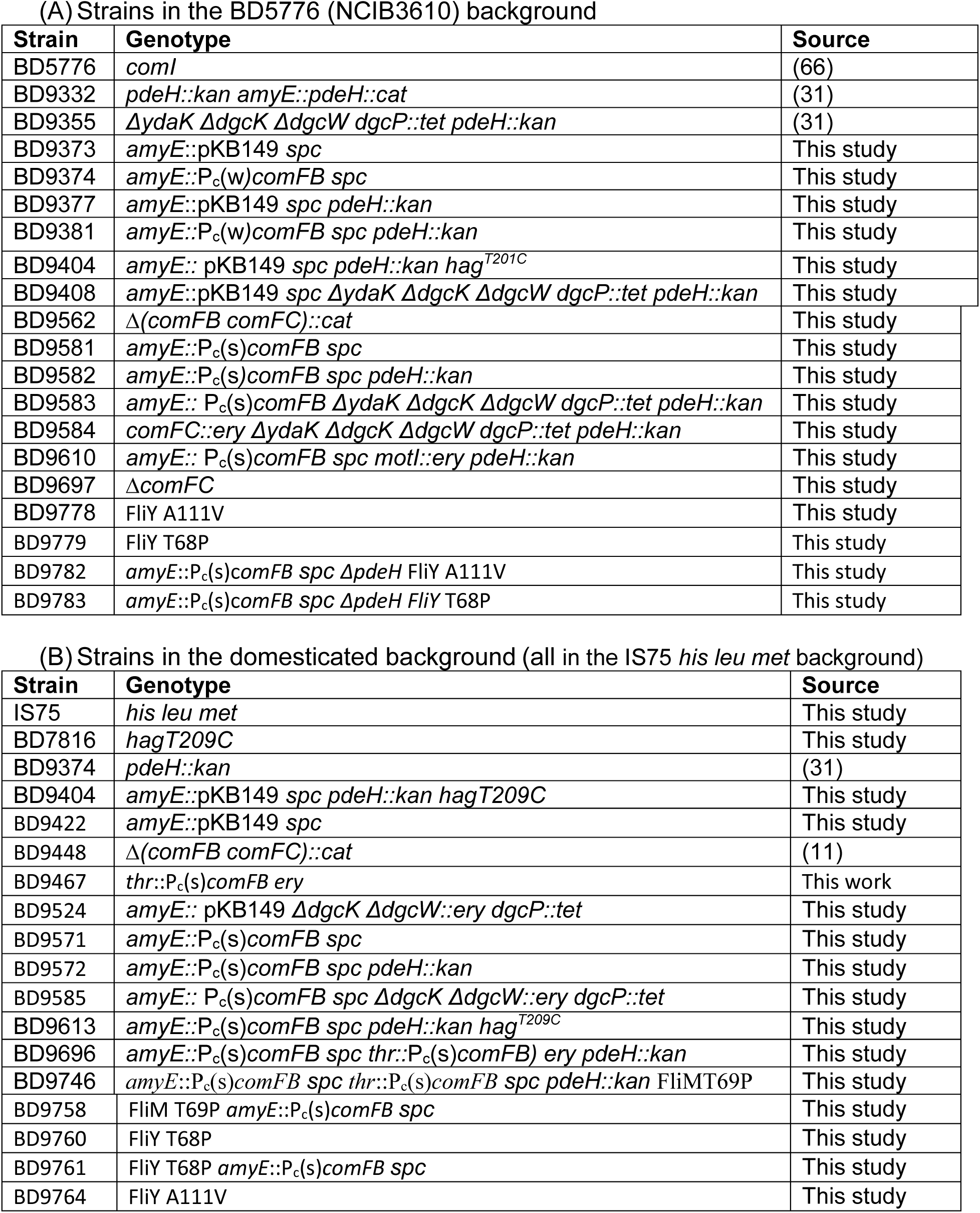
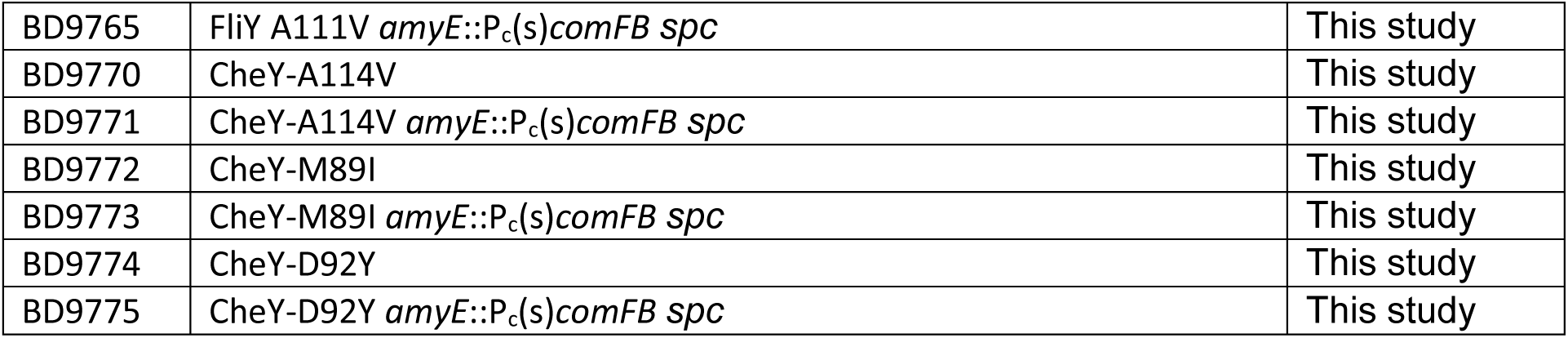
Strains.

(A) Strains in the BD5776 (NCIB3610) background
| Strain | Genotype | Source |
| --- | --- | --- |
| BD5776 | *comI* | (66) |
| BD9332 | *pdeH::kan amyE::pdeH::cat* | (31) |
| BD9355 | *ΔydaK ΔdgcK ΔdgcW dgcP::tet pdeH::kan* | (31) |
| BD9373 | *amyE::pKB149 spc* | This study |
| BD9374 | *amyE::P<sub>c</sub>(w)comFB spc* | This study |
| BD9377 | *amyE::pKB149 spc pdeH::kan* | This study |
| BD9381 | *amyE::P<sub>c</sub>(w)comFB spc pdeH::kan* | This study |
| BD9404 | *amyE:: pKB149 spc pdeH::kan hag<sup>T201C</sup>* | This study |
| BD9408 | *amyE::pKB149 spc ΔydaK ΔdgcK ΔdgcW dgcP::tet pdeH::kan* | This study |
| BD9562 | *Δ(comFB comFC)::cat* | This study |
| BD9581 | *amyE::P<sub>c</sub>(s)comFB spc* | This study |
| BD9582 | *amyE::P<sub>c</sub>(s)comFB spc pdeH::kan* | This study |
| BD9583 | *amyE:: P<sub>c</sub>(s)comFB ΔydaK ΔdgcK ΔdgcW dgcP::tet pdeH::kan* | This study |
| BD9584 | *comFC::ery ΔydaK ΔdgcK ΔdgcW dgcP::tet pdeH::kan* | This study |
| BD9610 | *amyE:: P<sub>c</sub>(s)comFB spc motl::ery pdeH::kan* | This study |
| BD9697 | *ΔcomFC* | This study |
| BD9778 | FliY A111V | This study |
| BD9779 | FliY T68P | This study |
| BD9782 | *amyE::P<sub>c</sub>(s)comFB spc ΔpdeH FliY A111V* | This study |
| BD9783 | *amyE::P<sub>c</sub>(s)comFB spc ΔpdeH FliY T68P* | This study |

(B) Strains in the domesticated background (all in the IS75 *his leu met* background)
| Strain | Genotype | Source |
| --- | --- | --- |
| IS75 | *his leu met* | This study |
| BD7816 | *hagT209C* | This study |
| BD9374 | *pdeH::kan* | (31) |
| BD9404 | *amyE::pKB149 spc pdeH::kan hagT209C* | This study |
| BD9422 | *amyE::pKB149 spc* | This study |
| BD9448 | *Δ(comFB comFC)::cat* | (11) |
| BD9467 | *thr::P<sub>c</sub>(s)comFB ery* | This work |
| BD9524 | *amyE:: pKB149 ΔdgcK ΔdgcW::ery dgcP::tet* | This study |
| BD9571 | *amyE::P<sub>c</sub>(s)comFB spc* | This study |
| BD9572 | *amyE::P<sub>c</sub>(s)comFB spc pdeH::kan* | This study |
| BD9585 | *amyE:: P<sub>c</sub>(s)comFB spc ΔdgcK ΔdgcW::ery dgcP::tet* | This study |
| BD9613 | *amyE::P<sub>c</sub>(s)comFB spc pdeH::kan hag<sup>T209C</sup>* | This study |
| BD9696 | *amyE::P<sub>c</sub>(s)comFB spc thr::P<sub>c</sub>(s)comFB ery pdeH::kan* | This study |
| BD9746 | *amyE::P<sub>c</sub>(s)comFB spc thr::P<sub>c</sub>(s)comFB spc pdeH::kan FliMT69P* | This study |
| BD9758 | FliM T69P *amyE::P<sub>c</sub>(s)comFB spc* | This study |
| BD9760 | FliY T68P | This study |
| BD9761 | FliY T68P *amyE::P<sub>c</sub>(s)comFB spc* | This study |
| BD9764 | FliY A111V | This study |
| BD9765 | FliY A111V *amyE*::P<sub>c</sub>(s)*comFB spc* | This study |
| BD9770 | CheY-A114V | This study |
| BD9771 | CheY-A114V *amyE*::P<sub>c</sub>(s)*comFB spc* | This study |
| BD9772 | CheY-M89I | This study |
| BD9773 | CheY-M89I *amyE*::P<sub>c</sub>(s)*comFB spc* | This study |
| BD9774 | CheY-D92Y | This study |
| BD9775 | CheY-D92Y *amyE*::P<sub>c</sub>(s)*comFB spc* | This study |

**Table 5.**
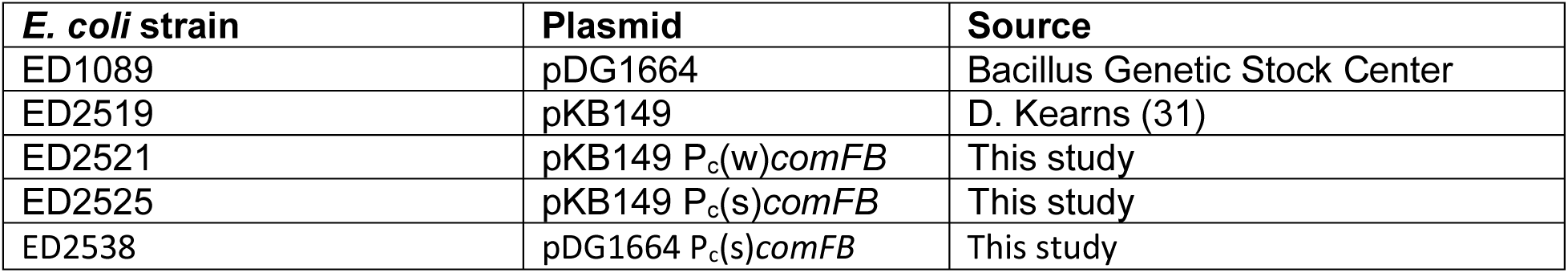
Plasmids.

Construction of the *ΔdgcK dgcW::erm dgcP::tet* knockout strain in the IS75 background was initiated by transformation with genomic DNA from DK392 (*dgcP::*tet), with selection for tetracycline. The Koo et al knockout collection (33) was used as a source of *dgcW::erm* and *dgcK::erm* deletions. *dgcK::erm* was first introduced into the *dgcP::tet* strain, and the erythromycin-resistance cassette was looped out using pDR244, as described (33). The resulting transformant was then transformed with *dgcW::erm* to produce the triple knockout erythromycin-and tetracycline-resistant strain (Table 4). For the insertion of constructs into *amyE*, plasmid DNA was linearized with *Sca*I and transformed into IS75 or BD5776, with selection for spectinomycin resistance. The resulting transformants were tested for insertion in *amyE* using starch plates stained with iodine. For insertion into *thrC*, plasmid DNA was digested with *Pvu*I and transformed into the appropriate *Bsu* strain, with erythromycin resistance selected, followed by tests for threonine auxotrophy and the absence of spectinomycin resistance.

### Molecular constructs

*amyE::Pc*(w)*FB*: The oligonucleotides ComFBWHinF (GCCATGAAGCTTCCGATGATGAGGAGCTG) and ComFBBamR (ATCGCCGGATCCGCGAATCACATAATAAACAGATC) were used in a PCR to isolate a *comFB* fragment containing the native RBS sequence that was then inserted between the *Hin*dIII and *Bam*HI sites of pKB149 (31). *amyE::Pc*(s)*FB*: The oligonucleotide ComFBSHinF (GCCATGAAGCTTAAAGGAGGAAACAATCATGCTTGTCAATTCAAAAGA) was used in a PCR with ComFBBamR to produce a *comFB* fragment containing the *gapA* RBS that was inserted between the *Hin*dIII and *Bam*HI sites of pKB149. *thrC*::Pc(s)*FB*: This construct was made using NEBuilder HiFi (New England Biolabs), and inserted into the *Hin*dIII site of pDG1664 for insertion in *thrC* (56). To obtain the Pc *gapA* RBS sequence, the oligonucleotides Pc(s)*F* (TGCTGCCTTCGGATCCTAGAAGCTTCTACACAGCCCAGTCCAG) and Pc(s)-R (CGTTTGACATGATTGTTTCCTCCTTTAAGCTTC) were used, with Pc(s)*FB* plasmid DNA as the template.

*ΔcomFC*: Strain BKE35450 (*ΔcomFC::erm*) (33) distributed by the Bacillus Genetic Stock Center was the source of this deletion. After transforming into the appropriate backgrounds, the Erm-resistance cassette with its promoter was removed, using plasmid pDR244, as described (33).

### Measurement of c-di-GMP binding to ComFB

*Bacillus subtilis* ComFB protein was heterologously expressed as an 8xHis-tagged protein in *E. coli* as described previously (7, 11). Briefly, the 8xHis-tagged constructs were expressed by overnight cultivation of *E. coli* cells at 20°C in the presence of 0.5 mM IPTG and purified by immobilized metal affinity chromatography using Ni^2+^-Sepharose resin (Cytiva^TM^), followed by size exclusion chromatography on a Superdex 200 Increase 10/300 GL column (GE HealthCare, Munich, Germany), as described previously (57). Protein purity was assessed by Coomassie-stained SDS-PAGE, and protein concentrations were determined using Bradford assay.

Binding of c-di-GMP to ComFB protein was analyzed by isothermal titration calorimetry (ITC), as described previously (58). In a buffer composed of (50 mM Tris/HCl, pH 8.0, 300 mM NaCl, 0.55 mM EDTA), the ITC measurements were conducted using a MicroCal PEAQ-ITC instrument (Malvern Panalytical, Westborough, MA, USA), at 25 °C, with a reference power of 10 μcal/s. Dissociation constants KD and ΔH values were calculated using the single binding site model with the MicroCal PEAQ-ITC Analysis Software (Malvern Panalytical). In the competition binding assays, the ComFB protein was incubated with 150 µM or 500 µM of (ATP, ADP, GTP, cAMP or cGMP) nucleotides and titrated against 1 mM of c-di-GMP.

Before ITC experiments were conducted, analytical size exclusion chromatography coupled to a multiangle light scattering (MALS) detector was carried out at room temperature, as described previously (57, 59, 60) using the ÄKTA purifier (GE Healthcare) on Superdex^TM^ 200 column. ASTRA software (Wyatt) was used for data analysis and molecular mass calculations using the MALS data. The elution volume was plotted against the UV signal and molecular mass.

### Swarm and Swim tests

Swarm expansion was measured using 0.7% LB agar as described by Gao *et al* (31). The plates were dried for about 10 minutes after spotting each strain in the center, then incubated at 37 °C in a humidified incubator with a water-filled tray on the bottom shelf. The diameter at the outermost edge of the swarm was measured at intervals. The swim assays were performed essentially as described by Hall et al (61). Petri dishes containing 0.3% LB agar were toothpick-inoculated with the desired strain and incubated overnight in a humidified incubator at 33 °C, then incubated at 37 °C. The plates were imaged in a ChemiDoc MP Bio-Rad imager when the fastest-swimming strain had nearly reached the perimeter of its plate.

### Western blotting

Anti-ComFB Western blotting from 12.5% tricine-SDS-polyacrylamide gels (62) was carried out by semidry blotting onto nitrocellulose membranes. The blots were developed using HRP-coupled goat anti-rabbit secondary antiserum (for ComFB and EFG) from Abcam and ECL kits from Amersham. The images were recorded with a Bio-Rad ChemiDoc MP imager. For loading controls, the top of the membrane was cut off after blotting and developed separately with anti-elongation factor G (anti-EFG) antiserum, a kind gift from Jonathan Dworkin (Columbia University Medical School). The anti-ComFB polyclonal antiserum was raised in rabbits by the ThermoFisher

Custom Antibody Service using ComFB protein purified as described by the Burton lab (11) from the His-tagged expression construct kindly provided by B. Burton.

### Flagellar labeling and microscopy

For fluorescence microscopy of flagella filaments, cells were grown at 37 °C in LB broth until an OD^600^ of 0.5–0.8 was reached. One milliliter of broth culture was harvested and resuspended in 50 μl of PBS containing 5 μg ml^-1^ Alexa Fluor 488 C5 maleimide (Thermofisher), and 1 μg/ml FM4-64. This suspension was incubated in the dark at room temperature for 5 min. Cells were then washed gently twice with 1 ml of PBS and resuspended to an OD^600^ of 10. Samples were observed by spotting 1 μl of the suspension on an agarose pad. Images were collected using a Nikon Eclipse Ti microscope equipped with an Orca Flash 4.0 Digital camera (Hamamatsu) and a Nikon TIRF 1.45 NA Plan Neofluor 100 oil-immersion objective. NIS-Elements AR (v 4.40, Nikon) software was used for image acquisition.

### Microscopic examination of swimming behavior

Cells were collected at the late logarithmic growth stage in LB, diluted, and placed in glass-bottom culture dishes (MatTek P35G-1.5-14-C) for observation. For observation of swimming in agar, the cells were briefly heated at 45 °C and mixed thoroughly with an equal volume of warm 0.6% agar before being placed in the dishes. For examination of cells from swarm plates, toothpicks were used to gently scrape the leading edges of swarms, and the cells were resuspended in LB for microscopic examination, as above. Videos were recorded using Nikon Elements with a 40X objective in a Nikon Ti microscope at 30 fps for 30 s. The .nd2 files were then exported in .mp4 format. The videos of suppressor strains were cropped to 0.25% and clipped to 10 s to reduce the file sizes, using iMovie. Videos of cells in soft agar were unclipped and uncropped, as were those for the analysis of swimming behavior.

#### Quantitative analysis of swimming behavior

To analyze bacterial swimming behavior, we developed a custom Python software package, MiMoSA (37) based on and extending the RABiTPy framework (38), an open-source Python package integrating image segmentation, particle tracking, and analysis of microbial motility. Briefly, videos or image sequences were acquired as described above, and cells were segmented from individual full-resolution frames using the integrated Omnipose framework with the bact_phase_affinity model (63). Trajectories were generated using a custom morphology- and motion-aware linker with a frame gap of 4, and subsequently filtered to exclude segmented objects inconsistent with individual cells, including debris, doublets, and cell chains. Trajectories were further manually curated to exclude those displaying circular or orbiting motility (39). For subsequent analysis, tracks were processed using a triangular smoothing filter; only consecutive-frame intervals were considered, and the first and final 10 frames of each trajectory were excluded. Swimming dynamics were analyzed using a velocity-based run-and-tumble detection method following Najafi et al. (39), based on the trajectory-based tumble-recognition framework developed by Masson et al. (64), and modified by Theves et al. (65). Tumbles were identified from temporally coincident local minima in translational speed, v(t), and local maxima in angular velocity magnitude, |ω(t)|. Persistence (*p*) is defined as cosθ, where θ is an angle that describes changes in direction during a run. R, cosΦ, is the equivalent metric for directional changesbetween runs.

More precisely, A speed minimum *v*(*t*_min_) was required to satisfy *Δv*⁄*v*(*t*_min_) ≥ 0.7, where *Δv* = *max*[*v*(*t*_1_) − *v*(*t*_min_), *v*(*t*_2_) − *v*(*t*_min_)] with the speed-defined tumble interval bounded by *v*(*t*) − *v*(*t*_min_) ≤ 0.2*Δv*. An angular maximum, *Δω* = *max*[|*ω*(*t*_max_)| − |*ω*(*t*_1_)|, |*ω*(*t*_max_)| − |*ω*(*t*_2_)|], was accepted if the magnitude of accumulated directional change angle, |*Δ*ф|, was greater than 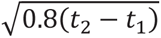 for that time interval. The directional change between successive runs was evaluated with the scalar of turning angle distribution *R*(ф), *R* = ⟨*cos* ф⟩, where ф is the inter-run turning angle as calculated using a four-point linear fit of the runs (39).

## ACKNOWLEDGEMENTS

This work was supported by NIH grants R01GM057720, awarded to DD, and R35-GM131783, awarded to DBK. RN was supported by NIH Grant R01GM136905 awarded to Carolyn Suzuki. The KAS laboratory is funded by grants from the German Research Foundation (DFG) as part of the Emmy Noether program (SE 3449/3-1), the priority research program (SPP2389; SE 3449/1-2), and by the collaborative research center SFB1381 (project number: 403222702; subproject B12). KAS also gratefully acknowledges the infrastructural support and funding from the collaborative research center MibiNet (SFB 1535 – Project ID 458090666) and the Cluster of Excellence “Controlling Microbes to Fight Infections (CMFI)” (EXC 2124 – 390838134). SS is funded by a scholarship from the Egyptian Ministry of Higher Education. We are also grateful to Filipp Oesterhelt (University of Tübingen) for facilitating access to the ITC.

## REFERENCES

1. Schavemaker PE, Lynch M. 2022. Flagellar energy costs across the tree of life. Elife https://www.ncbi.nlm.nih.gov/pubmed/35881430.

2. Jenal U, Reinders A, Lori C. 2017. Cyclic di-GMP: second messenger extraordinaire. Nat Rev Microbiol https://www.ncbi.nlm.nih.gov/pubmed/28163311:271-284.

3. Sun QX, Huang M, Zhang JY, Zeng X, Zhang CC. 2023. Control of Cell Size by c-di-GMP Requires a Two-Component Signaling System in the Cyanobacterium *Anabaena sp*. Strain PCC 7120. Microbiol Spectr https://www.ncbi.nlm.nih.gov/pubmed/36625639:e0422822.

4. Subramanian S, Gao X, Dann CE, 3rd, Kearns DB. 2017. MotI (DgrA) acts as a molecular clutch on the flagellar stator protein MotA in *Bacillus subtilis*. Proc Natl Acad Sci U S A https://www.ncbi.nlm.nih.gov/pubmed/29196522:13537-13542.

5. Zeng X, Huang M, Sun QX, Peng YJ, Xu X, Tang YB, Zhang JY, Yang Y, Zhang CC. 2023. A c-di-GMP binding effector controls cell size in a cyanobacterium. Proc Natl Acad Sci U S A https://www.ncbi.nlm.nih.gov/pubmed/36947515:e2221874120.

6. Wallner T, He C, Samir S, Lopes ES, Zeng X, Zhang CC, Selim KA, Yang Y, Wilde A. 2026. Regulation of cyanobacterial type IV pilus-dependent functions by interaction between a c-di-GMP receptor and two transcription factors. Microbiol Res https://www.ncbi.nlm.nih.gov/pubmed/42401049:128607.

7. Samir S, Elshereef AA, Alva V, Hahn J, Eck F, Celma L, Lopes ES, Thormann K, Dubnau D, Galperin MY, Selim KA. 2025. ComFB, a widespread family of c-di-NMP receptor proteins. Proc Natl Acad Sci U S A https://www.ncbi.nlm.nih.gov/pubmed/40966295:e2513041122.

8. Galperin MY, Chou SH. 2020. Structural Conservation and Diversity of PilZ-Related Domains. J Bacteriol 2020 Jan 29;202(4):e00664-19 https://www.ncbi.nlm.nih.gov/pubmed/31740493.

9. Samir S, Doello S, Enkerlin AM, Zimmer E, Haffner M, Muller T, Dengler L, Lambidis SP, Sivabalasarma S, Albers SV, Selim KA. 2025. The second messenger c-di-AMP controls natural competence via ComFB signaling protein. Cell Discov https://www.ncbi.nlm.nih.gov/pubmed/40745163:65.

10. Londono-Vallejo JA, Dubnau D. 1993. *comF*, a *Bacillus subtilis* late competence locus, encodes a protein similar to ATP-dependent RNA/DNA helicases. Mol Microbiol http://www.ncbi.nlm.nih.gov/htbin-post/Entrez/query?db=m&form=6&dopt=r&uid=0008412657:119-31.

11. Sysoeva TA, Bane LB, Xiao DY, Bose B, Chilton SS, Gaudet R, Burton BM. 2015. Structural characterization of the late competence protein ComFB from Bacillus subtilis. Biosci Rep 2015 Mar 31;35(2):e00183 https://www.ncbi.nlm.nih.gov/pubmed/25423369.

12. Diallo A, Foster HR, Gromek KA, Perry TN, Dujeancourt A, Krasteva PV, Gubellini F, Falbel TG, Burton BM, Fronzes R. 2017. Bacterial transformation: ComFA is a DNA-dependent ATPase that forms complexes with ComFC and DprA. Mol Microbiol https://www.ncbi.nlm.nih.gov/pubmed/28618091:741-754.

13. Dubnau D, Blokesch M. 2019. Mechanisms of DNA Uptake by Naturally Competent Bacteria. Annu Rev Genet https://www.ncbi.nlm.nih.gov/pubmed/31433955:217-237.

14. Maier B. 2020. Competence and Transformation in *Bacillus subtilis*. Curr Issues Mol Biol https://www.ncbi.nlm.nih.gov/pubmed/31950915:57-76.

15. Miras M, Dubnau D. 2016. A DegU-P and DegQ-Dependent Regulatory Pathway for the K-state in *Bacillus subtilis*. Front Microbiol http://www.ncbi.nlm.nih.gov/pubmed/27920766:1868.

16. Hahn J, Tanner AW, Carabetta VJ, Cristea IM, Dubnau D. 2015. ComGA-RelA interaction and persistence in the *Bacillus subtilis* K-state. Mol Microbiol http://www.ncbi.nlm.nih.gov/pubmed/25899641:454-71.

17. Haijema BJ, Hahn J, Haynes J, Dubnau D. 2001. A ComGA-dependent checkpoint limits growth during the escape from competence. Mol Microbiol:52–64.

18. Veening JW, Smits WK, Kuipers OP. 2008. Bistability, epigenetics, and bet-hedging in bacteria. Annu Rev Microbiol http://www.ncbi.nlm.nih.gov/entrez/query.fcgi?cmd=Retrieve&db=PubMed&dopt=Citation&list_uids=18537474:193-210.

19. Berka RM, Hahn J, Albano M, Draskovic I, Persuh M, Cui X, Sloma A, Widner W, Dubnau D. 2002. Microarray analysis of the *Bacillus subtilis* K-state: genome-wide expression changes dependent on ComK. Mol Microbiol http://www.ncbi.nlm.nih.gov/entrez/query.fcgi?cmd=Retrieve&db=PubMed&dopt=Citation&list_uids=11918817:1331–45.

20. Hamoen LW, Smits WK, de Jong A, Holsappel S, Kuipers OP. 2002. Improving the predictive value of the competence transcription factor (ComK) binding site in *Bacillus subtilis* using a genomic approach. Nucleic Acids Res http://www.ncbi.nlm.nih.gov/entrez/query.fcgi?cmd=Retrieve&db=PubMed&dopt=Citation&list_uids=12490720:5517-28.

21. Ogura M, Yamaguchi H, Kobayashi K, Ogasawara N, Fujita Y, Tanaka T. 2002. Whole-genome analysis of genes regulated by the *Bacillus subtilis* competence transcription factor ComK. J Bacteriol http://www.ncbi.nlm.nih.gov/entrez/query.fcgi?cmd=Retrieve&db=PubMed&dopt=Citation&list_uids=11948146:2344-51.

22. Hernandez-Plaza A, Szklarczyk D, Botas J, Cantalapiedra CP, Giner-Lamia J, Mende DR, Kirsch R, Rattei T, Letunic I, Jensen LJ, Bork P, von Mering C, Huerta-Cepas J. 2023. eggNOG 6.0: enabling comparative genomics across 12 535 organisms. Nucleic Acids Res https://www.ncbi.nlm.nih.gov/pubmed/36399505:D389-D394.

23. Mohari B, Licata NA, Kysela DT, Merritt PM, Mukhopadhay S, Brun YV, Setayeshgar S, Fuqua C. 2015. Novel pseudotaxis mechanisms improve migration of straight-swimming bacterial mutants through a porous environment. mBio https://www.ncbi.nlm.nih.gov/pubmed/25714707:e00005.

24. Wolfe AJ, Berg HC. 1989. Migration of bacteria in semisolid agar. Proc Natl Acad Sci U S A https://www.ncbi.nlm.nih.gov/pubmed/2674941:6973-7.

25. Chou SH, Galperin MY. 2016. Diversity of Cyclic Di-GMP-Binding Proteins and Mechanisms. J Bacteriol https://www.ncbi.nlm.nih.gov/pubmed/26055114:32-46.

26. Herzberg C, Meissner J, Warneke R, Stulke J. 2023. The many roles of cyclic di-AMP to control the physiology of *Bacillus subtilis*. Microlife https://www.ncbi.nlm.nih.gov/pubmed/37954098:uqad043.

27. Mantovani O, Haffner M, Selim KA, Hagemann M, Forchhammer K. 2023. Roles of second messengers in the regulation of cyanobacterial physiology: the carbon-concentrating mechanism and beyond. Microlife https://www.ncbi.nlm.nih.gov/pubmed/37223741:uqad008.

28. Hengge R. 2009. Principles of c-di-GMP signalling in bacteria. Nat Rev Microbiol https://www.ncbi.nlm.nih.gov/pubmed/19287449:263-73.

29. Khan F, Jeong GJ, Tabassum N, Kim YM. 2023. Functional diversity of c-di-GMP receptors in prokaryotic and eukaryotic systems. Cell Commun Signal https://www.ncbi.nlm.nih.gov/pubmed/37749602:259.

30. Kearns DB, Losick R. 2003. Swarming motility in undomesticated *Bacillus subtilis*. Mol Microbiol https://www.ncbi.nlm.nih.gov/pubmed/12864845:581-90.

31. Gao X, Mukherjee S, Matthews PM, Hammad LA, Kearns DB, Dann CE, 3rd. 2013. Functional characterization of core components of the *Bacillus subtilis* cyclic-di-GMP signaling pathway. J Bacteriol https://www.ncbi.nlm.nih.gov/pubmed/23893111:4782-92.

32. Chen Y, Chai Y, Guo JH, Losick R. 2012. Evidence for cyclic Di-GMP-mediated signaling in *Bacillus subtilis*. J Bacteriol https://www.ncbi.nlm.nih.gov/pubmed/22821967:5080-90.

33. Koo BM, Kritikos G, Farelli JD, Todor H, Tong K, Kimsey H, Wapinski I, Galardini M, Cabal A, Peters JM, Hachmann AB, Rudner DZ, Allen KN, Typas A, Gross CA. 2017. Construction and Analysis of Two Genome-Scale Deletion Libraries for *Bacillus subtilis*. Cell Syst https://www.ncbi.nlm.nih.gov/pubmed/28189581:291-305 e7.

34. Liu J, Zuber P. 1998. A molecular switch controlling competence and motility: competence regulatory factors ComS, MecA, and ComK control sigmaD-dependent gene expression in *Bacillus subtilis*. J Bacteriol http://www.ncbi.nlm.nih.gov/cgi-bin/Entrez/referer?http://jb.asm.org/cgi/content/full/180/16/4243:4243-51.

35. Blair KM, Turner L, Winkelman JT, Berg HC, Kearns DB. 2008. A molecular clutch disables flagella in the *Bacillus subtilis* biofilm. Science http://www.ncbi.nlm.nih.gov/entrez/query.fcgi?cmd=Retrieve&db=PubMed&dopt=Citation&list_uids=18566286:1636-8.

36. Bhattacharjee T, Datta SS. 2019. Bacterial hopping and trapping in porous media. Nat Commun https://www.ncbi.nlm.nih.gov/pubmed/31061418:2075.

37. Noland R. 2026. MiMoSA [Computer software], https://github.com/rnprgh/MiMoSA

38. Sen S, Vairagare I, Gosai J, Shrivastava A. 2025. RABiTPy: an open-source Python software for rapid, AI-powered bacterial tracking and analysis. BMC Bioinformatics https://www.ncbi.nlm.nih.gov/pubmed/40383775:127.

39. Najafi J, Shaebani MR, John T, Altegoer F, Bange G, Wagner C. 2018. Flagellar number governs bacterial spreading and transport efficiency. Sci Adv https://www.ncbi.nlm.nih.gov/pubmed/30263953:eaar6425.

40. Partridge JD, Nhu NTQ, Dufour YS, Harshey RM. 2019. Escherichia coli Remodels the Chemotaxis Pathway for Swarming. mBio https://www.ncbi.nlm.nih.gov/pubmed/30890609.

41. Partridge JD, Nhu NTQ, Dufour YS, Harshey RM. 2020. Tumble Suppression Is a Conserved Feature of Swarming Motility. mBio 2020 Jun 16;11(3):e01189-20 https://www.ncbi.nlm.nih.gov/pubmed/32546625.

42. Chandrangsu P, Helmann JD. 2016. Intracellular Zn(II) Intoxication Leads to Dysregulation of the PerR Regulon Resulting in Heme Toxicity in *Bacillus subtilis*. PLoS Genet https://www.ncbi.nlm.nih.gov/pubmed/27935957:e1006515.

43. Bedrunka P, Graumann PL. 2017. Subcellular clustering of a putative c-di-GMP-dependent exopolysaccharide machinery affecting macro colony architecture in *Bacillus subtilis*. Environ Microbiol Rep https://www.ncbi.nlm.nih.gov/pubmed/27897378:211-222.

44. Guttenplan SB, Blair KM, Kearns DB. 2010. The EpsE flagellar clutch is bifunctional and synergizes with EPS biosynthesis to promote *Bacillus subtilis* biofilm formation. PLoS Genet https://www.ncbi.nlm.nih.gov/pubmed/21170308:e1001243.

45. Paul K, Nieto V, Carlquist WC, Blair DF, Harshey RM. 2010. The c-di-GMP binding protein YcgR controls flagellar motor direction and speed to affect chemotaxis by a "backstop brake" mechanism. Mol Cell https://www.ncbi.nlm.nih.gov/pubmed/20346719:128-39.

46. Fang X, Gomelsky M. 2010. A post-translational, c-di-GMP-dependent mechanism regulating flagellar motility. Mol Microbiol https://www.ncbi.nlm.nih.gov/pubmed/20444091:1295-305.

47. Hou YJ, Yang WS, Hong Y, Zhang Y, Wang DC, Li DF. 2020. Structural insights into the mechanism of c-di-GMP-bound YcgR regulating flagellar motility in Escherichia coli. J Biol Chem https://www.ncbi.nlm.nih.gov/pubmed/31836667:808-821.

48. Han Q, Wang SF, Qian XX, Guo L, Shi YF, He R, Yuan JH, Hou YJ, Li DF. 2023. Flagellar brake protein YcgR interacts with motor proteins MotA and FliG to regulate the flagellar rotation speed and direction. Front Microbiol https://www.ncbi.nlm.nih.gov/pubmed/37125196:1159974.

49. Ha DG, O’Toole GA. 2015. c-di-GMP and its Effects on Biofilm Formation and Dispersion: a Pseudomonas Aeruginosa Review. Microbiol Spectr https://www.ncbi.nlm.nih.gov/pubmed/26104694:MB-0003-2014.

50. Chen X, Wang L, Sourjik V. 2025. Swimming or sessile: the interplay between c-di-GMP signalling and flagellar motility. Curr Opin Microbiol https://www.ncbi.nlm.nih.gov/pubmed/40638951:102632.

51. Diethmaier C, Chawla R, Canzoneri A, Kearns DB, Lele PP, Dubnau D. 2017. Viscous drag on the flagellum activates *Bacillus subtilis* entry into the K-state. Mol Microbiol https://www.ncbi.nlm.nih.gov/pubmed/28800172:367-380.

52. Holscher T, Schiklang T, Dragos A, Dietel AK, Kost C, Kovacs AT. 2018. Impaired competence in flagellar mutants of *Bacillus subtilis* is connected to the regulatory network governed by DegU. Environ Microbiol Rep https://www.ncbi.nlm.nih.gov/pubmed/29124898:23-32.

53. Mukherjee S, Kearns DB. 2014. The structure and regulation of flagella in *Bacillus subtilis*. Annu Rev Genet https://www.ncbi.nlm.nih.gov/pubmed/25251856:319-40.

54. Dubnau D, Davidoff-Abelson R. 1971. Fate of transforming DNA following uptake by competent *Bacillus subtilis*. I. Formation and properties of the donor-recipient complex. J Mol Biol:209–221.

55. Patrick JE, Kearns DB. 2008. MinJ (YvjD) is a topological determinant of cell division in *Bacillus subtilis*. Mol Microbiol http://www.ncbi.nlm.nih.gov/entrez/query.fcgi?cmd=Retrieve&db=PubMed&dopt=Citation&list_uids=18976281:1166-79.

56. Guerout-Fleury AM, Frandsen N, Stragier P. 1996. Plasmids for ectopic integration in *Bacillus subtilis*. Gene http://www.ncbi.nlm.nih.gov/entrez/query.fcgi?cmd=Retrieve&db=PubMed&dopt=Citation&list_uids=8973347:57-61.

57. Selim KA, Lapina T, Forchhammer K, Ermilova E. 2020. Interaction of N-acetyl-l-glutamate kinase with the PII signal transducer in the non-photosynthetic alga *Polytomella parva*: Co-evolution towards a hetero-oligomeric enzyme. FEBS J https://www.ncbi.nlm.nih.gov/pubmed/31287617:465-482.

58. Selim KA, Haase F, Hartmann MD, Hagemann M, Forchhammer K. 2018. P(II)-like signaling protein SbtB links cAMP sensing with cyanobacterial inorganic carbon response. Proc Natl Acad Sci USA https://www.ncbi.nlm.nih.gov/pubmed/29735650:E4861-E4869.

59. Selim KA, Haffner M, Watzer B, Forchhammer K. 2019. Tuning the in vitro sensing and signaling properties of cyanobacterial PII protein by mutation of key residues. Sci Rep https://www.ncbi.nlm.nih.gov/pubmed/31831819:18985.

60. Walter J, Selim KA, Leganes F, Fernandez-Pinas F, Vothknecht UC, Forchhammer K, Aro EM, Gollan PJ. 2019. A novel Ca(2+)-binding protein influences photosynthetic electron transport in *Anabaena sp*. PCC 7120. Biochim Biophys Acta Bioenerg https://www.ncbi.nlm.nih.gov/pubmed/31034800:519-532.

61. Hall AN, Subramanian S, Oshiro RT, Canzoneri AK, Kearns DB. 2018. SwrD (YlzI) Promotes Swarming in *Bacillus subtilis* by Increasing Power to Flagellar Motors. J Bacteriol https://www.ncbi.nlm.nih.gov/pubmed/29061663.

62. Schagger H, von Jagow G. 1987. Tricine-sodium dodecyl sulfate-polyacrylamide gel electrophoresis for the separation of proteins in the range from 1 to 100 kDa. Anal Biochem:368–379.

63. Cutler KJ, Stringer C, Lo TW, Rappez L, Stroustrup N, Brook Peterson S, Wiggins PA, Mougous JD. 2022. Omnipose: a high-precision morphology-independent solution for bacterial cell segmentation. Nat Methods https://www.ncbi.nlm.nih.gov/pubmed/36253643:1438-1448.

64. Masson JB, Voisinne G, Wong-Ng J, Celani A, Vergassola M. 2012. Noninvasive inference of the molecular chemotactic response using bacterial trajectories. Proc Natl Acad Sci U S A https://www.ncbi.nlm.nih.gov/pubmed/22307649:1802-7.

65. Theves M, Taktikos J, Zaburdaev V, Stark H, Beta C. 2013. A bacterial swimmer with two alternating speeds of propagation. Biophys J https://www.ncbi.nlm.nih.gov/pubmed/24138867:1915-24.

66. Konkol MA, Blair KM, Kearns DB. 2013. Plasmid-encoded ComI Inhibits competence in the ancestral 3610 strain of *Bacillus subtilis*. J Bacteriol http://www.ncbi.nlm.nih.gov/pubmed/23836866:4085-93.

