## Supplement Figures S1 and S2 for "ComFB bound to c-di-GMP reduces swimming speed and tumbling frequency in *B. subtilis*"

**Figure S1**

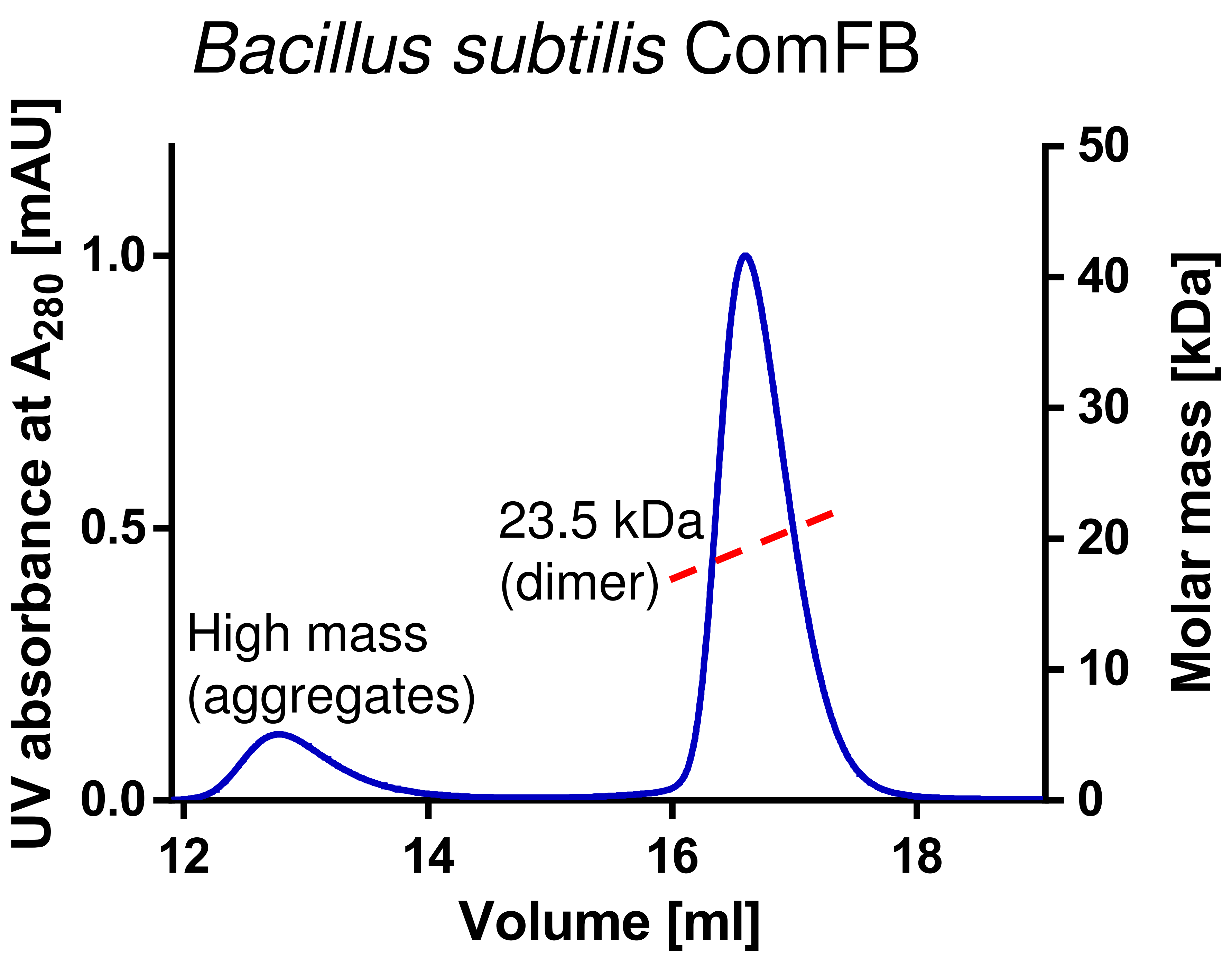


Figure S1. Oligomeric state of FB protein as determined by size exclusion chromatography coupled to multiangle light scattering (SEC-MALS). SEC-MALS showed a species distribution of ~ 23.5 kDa for FB protein, indicating that FB (the theoretical molecular mass of a monomer with an 8xHis tag is 12.1 kDa) is a dimer in solution.

### Figure S2

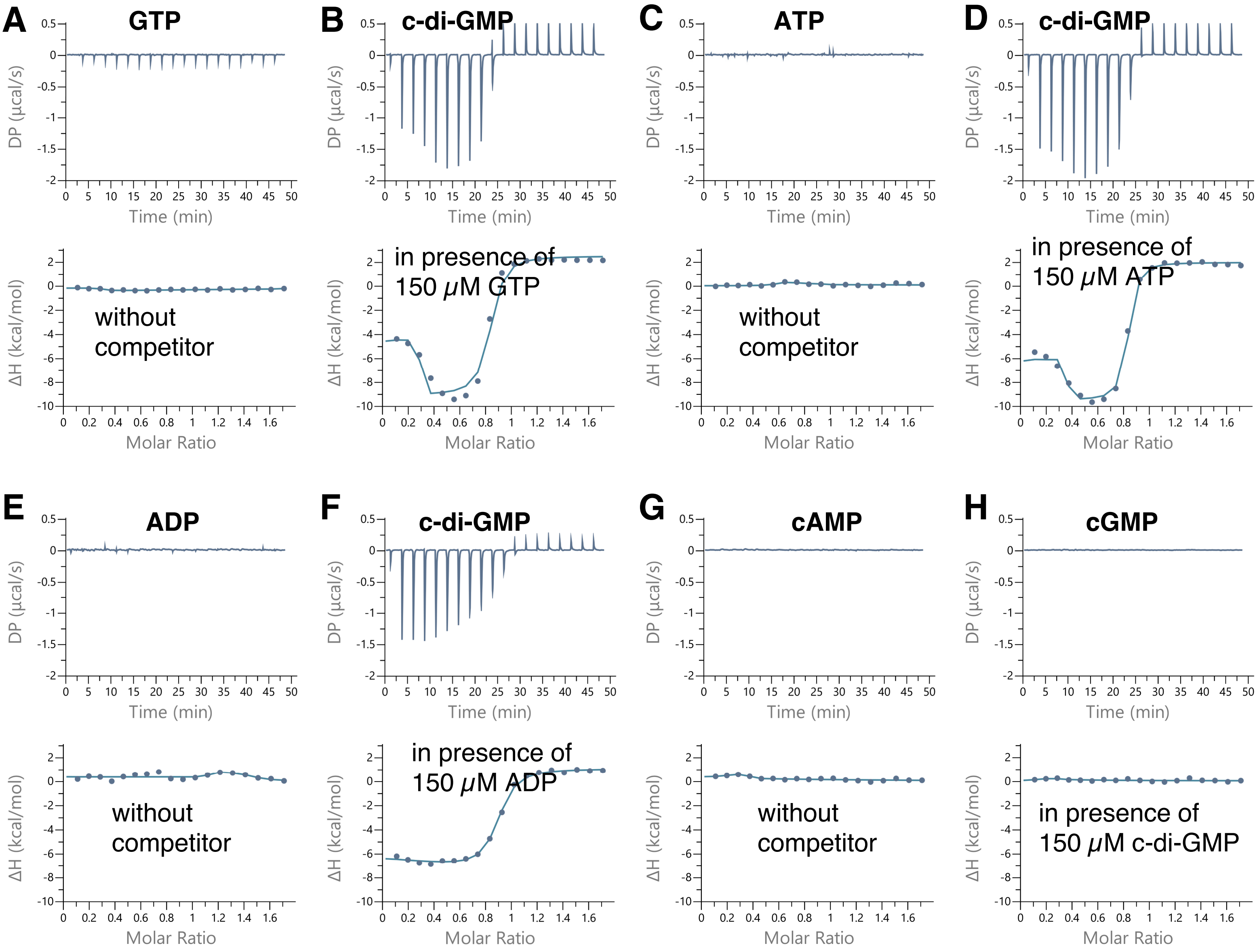


Figure S2. Isothermal titration calorimetry (ITC) analysis of GTP, ATP, ADP, cAMP, cGMP, and c-di-GMP binding to FB, as indicated. Upper panels show the raw ITC data in the form of heat produced during the titration of the respective nucleotide on FB protein; lower panels show the binding isotherms and the best-fit curves according to the one binding site model. In case of c-di-GMP titrations, the FB protein was first incubated with 150 μM of GTP, ATP, ADP, cAMP, or cGMP, as indicated.

Supplemental Figure Legends

Figure S1. Oligomeric state of FB protein as determined by size exclusion chromatography coupled to multiangle light scattering (SEC-MALS). SEC-MALS showed a species distribution of ~ 23.5 kDa for FB protein, indicating that FB (the theoretical molecular mass of a monomer with an 8xHis tag (12.1 kDa) is a dimer in solution.

Figure S2. Isothermal titration calorimetry (ITC) analysis of GTP, ATP, ADP, cAMP, cGMP, and c-di-GMP binding to FB, as indicated. Upper panels show the raw ITC data in the form of heat produced during the titration of the respective nucleotide on FB protein; lower panels show the binding isotherms and the best-fit curves according to the one-binding-site model. In case of c-di GMP titrations, the FB protein was first incubated with 150 µM of GTP, ATP, ADP, cAMP, or cGMP, as indicated.
